# Decoupled spectral tuning and eye size diversification patterns in an Antarctic adaptive radiation

**DOI:** 10.1101/2022.09.28.509872

**Authors:** Ella B. Yoder, Elyse Parker, Alexandra Tew, Christopher D. Jones, Alex Dornburg

**Affiliations:** Department of Bioinformatics and Genomics, University of North Carolina at Charlotte, Charlotte, NC USA; Research Triangle High School, Durham, NC USA; Department of Ecology & Evolutionary Biology, Yale University, New Haven, CT 06520, USA; Antarctic Ecosystem Research Division, NOAA Southwest Fisheries Science Center, La Jolla, CA 92037, USA

**Keywords:** opsin, tuning site, icefish, notothenioids

## Abstract

Evolutionary transitions in water column usage have played a major role in shaping ray-finned fish diversity. However, the extent to which vision-associated trait complexity and water column usage is coupled remains unclear. Here we investigate the relationship between depth niche, eye size, and the molecular basis of light detection across the Antarctic notothenioid adaptive radiation. Using a phylogenetic comparative framework, we integrate sequence analyses of opsin tuning sites with data on eye size and depth occupancy from over two decades of NOAA trawl-based surveys. We find a consistent signature of changes in tuning sites suggestive of shifts in their ability to detect lower wavelengths of light. These represent repeated instances of independent tuning site changes across the notothenioid phylogeny that are generally not associated with habitat depth or species eye size. We further reveal an acceleration in the rate of eye size diversification nearly 20 million years after the initial radiation that has manifested in high levels of eye size divergence among closely related taxa. Collectively, our results strongly support a decoupling of the diversification dynamics between opsin tuning sites, eye size and depth, providing a new perspective of the evolution of the visual system in this iconic adaptive radiation.

## Introduction

Evolutionary changes in water column usage have shaped the diversification of aquatic organisms across the Tree of Life (Rosa et al. 2008; Modica et al. 2020). This is particularly the case in ray-finned fishes (Actinopterygii) where transitions between benthic and pelagic habitat use or depth usage have been repeatedly linked to the diversification of morphological (Smith and Brown 2002; Ingram 2011; John et al. 2022), physiological (Brown and Thatje 2014), or ecological traits (Fukunaga et al. 2016; Costello and Chaudhary 2017). While it is clear that stratification of the water column has played a pivotal role in shaping the evolution of traits such as those associated with feeding (Drazen and Sutton 2017) or locomotion (Martinez et al. 2021), the evolution of the visual system has received comparatively less attention.

Recent investigations of individual marine fishes have highlighted that changes in water column usage are often associated with extremes in vision. In some taxa the perceivable visual spectrum may become reduced to a fraction of that detectable by closely related taxa (Marshall et al. 2015). In these cases, color vision is entirely lost (monochromacy) or reduced to two channels (dichromacy) that function to discriminate objects against a background color (Lythgoe et al. 1994; Marshall et al. 2015). Conversely, numerous lineages utilize three (trichromacy), four (tetrachromacy), or more spectral channels, thereby altering their utilization of a given light environment and providing potential advantages to signaling or color discrimination (Sabbah et al. 2013; Stieb et al. 2017; Marshall et al. 2019). Additionally, eye size can be extremely heterogeneous for taxa in dim-light environments. Large expansions in eye size capture more light that can signal approaching predators or aid in prey detection (Kotrschal et al. 2017; Vinterstare et al. 2020). Alternatively, reductions in eye size may reflect organisms minimizing a trade-off in neural investment between photo and chemo-perception (Iglesias et al. 2018). To develop general rules that gave rise to these patterns of vision-associated trait complexity and reduction, investigations of clades radiating across the water-column in different environments are critically needed.

Antarctic notothenioids represent an exemplar system for investigating the diversification of the visual system across water-column niches. These fishes comprise the bulk of the near-shore fish diversity of the Antarctic and are a critical food source for many Antarctic species (penguins, seals, whales) (La Mesa et al. 2004b), the basis of a multi-billion dollar fishing industry (Constable et al. 2000; Abrams 2014), and one of few examples of adaptive radiation in a marine environment (Eastman 2000; Matschiner et al. 2011; Daane et al. 2019). Moreover, their ability to rapidly diversify into water column niches that span hundreds of meters forms a core ecological basis of this radiation (Rutschmann et al. 2011; Near et al. 2012; Parker et al. 2022). The rapid diversification of water-column niches has occurred in part due to the ecological opportunities generated by cyclical ice scour events that obliterate entire benthic communities across thousands of kilometers of the Antarctic continental shelf (Thatje et al. 2005, 2008; Near et al. 2012; Strugnell et al. 2012; Dornburg et al. 2017). Following the decimation of the incumbent fauna, previously occupied niches are emptied and provide new ecological opportunities for surviving lineages. This cyclical pattern of extinction and speciation has resulted in evolutionary close relatives often being highly divergent in their water-column niche (Parker et al. 2022). Given the changes in light attenuation associated with changes in depth, how the diversification across the water column has shaped the evolution of the notothenioid visual system represents an important aspect of this adaptive radiation that can now be illuminated by the combination of recently collected molecular data and decades of sampling efforts.

At the molecular level, the process of vision is initiated through a light induced conformational change in opsins, photopigments which are bound to a vitamin A-derived chromophore (Terakita 2005). Phototransduction occurs through the interaction between opsin protein and the chromophore which determines the maximum spectrum absorbance (λ_max_) of the opsin. A number of residues have been identified in opsins as “tuning sites” that influence which wavelengths (λ_max_) trigger phototransduction and the wavelengths of detectable light (Musilova et al. 2021). Across vertebrates, rods contain only one class of opsin (rhodopsin-like-1; Rh1) that is responsible for scotopic (dim-light) vision (Pisani et al. 2006; Yokoyama et al. 2008). Cones contain four classes of opsins that are responsible for photopic (bright-light) vision. Short-wavelength-sensitive-1 (SWS1) and short-wavelength-sensitive-2 (SWS2) absorb in the ultraviolet [UV; peak spectral sensitivity (λ_max_)=355–450 nm] and violet/blue (λ_max_=415–490 nm) regions of the spectrum, respectively. Middle-wavelength-sensitive / rhodopsin-like-2 (Rh2) is most sensitive to the central (green) waveband (λ_max_=470–535 nm), and long wavelength sensitive (LWS) is tuned towards the spectrum’s red end (λ_max_=490–570 nm) (Yokoyama 2002). Early investigations of notothenioid opsins suggested that these lineages possess a limited repertoire of opsins relative to other ray-finned fishes (Pointer et al. 2005). While a reduction of opsins could be expected for organisms living with seasonal light availability at varying depths, it is now clear that this is not the case. Instead, notothenioid opsin diversity is on par with that of other fishes possessing complex color tetrachromatic vision (Hunt et al. 2001; Rennison et al. 2012; Lin et al. 2017; Musilova et al. 2021). The finding of possible tetrachromatic vision raises the question of how opsin tuning sites evolved during the radiation of this clade, and whether changes in tuning sites are associated with changes in eye size.

As a predominately polar radiation, notothenioids experience extreme seasonal differences in sunlight modulated by the tilt of the earth. Additionally, many notothenioids live at or below the limits of the “photic zone” (150-220 m deep), or below layers of surface ice and snow that inhibit sunlight transmittance at around 500 nm with substantially reduced intensity (Pointer et al. 2005). Although notothenioids are all subjected to dim-light conditions, notothenioid eye sizes vary dramatically and encompass a level of diversity as high as that exhibited collectively across all the near-shore coastal fishes of New Zealand (Montgomery and Macdonald 1998). The mechanisms promoting the diversification of eye sizes in notothenioids remain unexplored as does the evolutionary relationship between eye size and opsin tuning sites. On the one hand, eye size may be modulated by changes in depth usage and associated with specific tuning site changes towards the detection of shorter wavelengths. Such a hypothesis would be in line with classic expectations of strong selection by the photic environment and associated by convergence in eye size and tuning sites across the notothenioid phylogeny. On the other hand, eye size may not be a labile trait over short evolutionary timescales (de Busserolles et al. 2013) and could be decoupled from changes in tuning sites occurring as lineages transition to alternate water-column niches. In this case, eye size would exhibit a signature of conservation within clades converging on specific tuning sites. Determining the extent to which these types of broad hypotheses explain the diversification of opsin tuning site and eye size evolution in notothenioids represents a pivotal step in developing an understanding of the role of the visual system in this adaptive radiation. Moreover, forecasts of Antarctica’s changing climate make such insights timely for better understanding the stewardship needs of these vital living marine resources (Patarnello et al. 2011; Mintenbeck and Torres 2017).

Here we investigate the evolutionary history of opsin tuning sites and eye size across notothenioids using a time calibrated phylogenetic framework. To assess evidence of tuning site changes that equate optic adaptations at the molecular level, we first accessed publicly available genomes, transcriptomes, and additional sequences to identify and analyze opsins from all major notothenioid lineages. We then assessed the relationship between tuning site changes and depth by coupling these analyses with data on the depth distributions of notothenioids using over two decades of NOAA trawl-based surveys. Finally, we tested the extent to which the evolution of changes in tuning sites, water column usage, and relative eye size were coupled using a series of phylogenetic comparative analyses. Collectively, these results reveal unusual patterns of water-column and vision-associated diversification that strongly support a decoupling of the diversification dynamics of these traits.

## Methods

### Identification and comparison of notothenioid opsin sequences

Zebrafish opsin sequences LWS1 (BAC24127.1), LWS2 (BAC24128.1), Rh2-1 (BAC24129.1), Rh2-2 (BAC24130.1), Rh2-3 (BAC24131.1), Rh2-4 (BAC24132.1), SWS1 (BAC24134.1), SWS2 (BAC24133.1), and Rh1-1 (NP_571159.1) and Rh1-2 (BAC21668.1) (Hamaoka et al. 2002; Chinen et al. 2003; Morrow et al. 2011, 2017) were used as queries for BLASTp searches restricted to Notothenioidei (taxid: 8205), giant grouper (*Epinephelus lanceolatus* - taxid: 310571), and wolf eel (*Anarrhichthys ocellatus* - taxid: 433405). We additionally queried available notothenioid genomes on NCBI and the *Eleginops maclovinus* genome (Chen et al. 2019) available through GIGADB (DOI: 10.5524/102163).

Sequences were identified using HMMER (Eddy 2009) with an e-value threshold of 10 x e^-6^. Training profiles for HMMer were built using a fasta alignment of known opsin receptors from NCBI identified above. Candidate opsins were aligned to the reference sequence dataset for manual inspection. All protein sequences were aligned using MAFFT (Katoh 2002) and inspected via Aliview (Larsson 2014). As several available notothenioid genomes were sequenced with low (10x) coverage, instances of fragmented opsin genes within scaffolds were excluded from analysis as meaningful tuning site comparisons would not be possible. Additionally, non-visual extra-ocular rhodopsin, (exo-rh1) sequences, which are similar to Rh1 but expressed in the pineal gland (Chen et al. 2018; Fujiyabu et al. 2019), were excluded from this study.

Opsin protein sequences were imported into Geneious Prime 2021.2 (https://www.geneious.com) where they were aligned using the Clustal Omega (Sievers and Higgins 2014) plugin in Geneious Prime with no adjustments. The identity of opsins were assessed using maximum likelihood based phylogenetic inference in IQTree2 (Minh et al. 2020) conditioned on the best fit model of amino acid substitution with the candidate pool of substitution rates spanning all common amino acid exchange rate matrices (JTT, WAG, etc). The candidate pool of substitution models additionally included protein mixture models such as empirical profile mixture models (Quang et al. 2008; Minh et al. 2021), as well as parameters to accommodate among-site rate variation (discrete gamma or free rate model). The best fit model for each alignment was selected using Bayesian information criterion with node support assessed via 1,000 ultrafast bootstrap replicates.

### Collection of Ecological and Morphological Data

To complement our investigation of opsin tuning sites across the notothenioid radiation, we assessed patterns of diversification in notothenioid eye size and investigated the extent to which evolutionary changes in eye size are modulated by diversification along the water column. We used the program TpsDIG2 (Rohlf 2006) to quantify eye size and body size from digital images of 161 specimens representing 61 notothenioid species (Supplemental Table S1). Where possible, we sampled three individuals per species, but in some cases only one or two specimens were available. Eye diameter was measured from the most anterior point of the eye to the most posterior point of the eye. Body size was represented using Standard Length (SL), measured as the distance from the tip of the snout to the posterior end of the hypural plate. We additionally measured maximum body depth as the maximum distance between the outer dorsal and outer ventral surfaces and head length as the distance from the tip of the snout to the posterior edge of the operculum. We then compiled data on water column usage for the same species measured in our morphological dataset. Depth occupancy was represented using three different measures: (1) mean depth of capture for 44 of our 61 focal species calculated from over 19,900 trawl records of adult specimens from the U.S. Antarctic Marine Resources Program (U.S. AMLR) Antarctic finfish surveys between 1998 and 2018; (2) minimum reported depth occupancy for all 61 focal species (summarized in (Eastman 2017)); and (3) maximum reported depth occupancy for all 61 focal species (summarized in (Eastman 2017)). For 46 of our focal species, we represented water column niche using data on mean percentage buoyancy (%B), which is a measure of the overall body density of species calculated as the percentage of body weight in air supported in seawater (Eastman and DeVries 1982; Near et al. 2012; Eastman 2020).

### Calculating the impact of water column usage and buoyancy on eye size

We used time-calibrated phylogenies taken from Parker et al. (Parker et al. 2022) to conduct a series of comparative phylogenetic analyses that evaluated the relationships among evolutionary changes in water column usage and diversification of eye size in notothenioids. Measurements of morphological traits are expected to scale with measures of body size (Berner 2011), and eye diameter is correlated with body length, body depth, or head length in notothenioids. As such, we represented eye size using residuals from a phylogenetic generalized least squares (PGLS) regression [implemented using the gls function in the nlme R package ((Pinheiro et al. n.d.))] of eye diameter on head length. This approach enables us to fit linear models of trait correlation while also accounting for the covariance among traits due to phylogeny. For all tests, we used the corBrownian function in the ape R package ((Paradis et al. 2004)) to define the trait covariance structure using a Brownian Motion (BM) model of trait evolution on a time calibrated notothenioid phylogeny derived from Near et al. (Near et al. 2018). We then used PGLS regression under a BM model of evolution to test for correlations between residual eye size and depth occupancy (represented as mean depth of capture, minimum reported depth, or maximum reported depth) as well as between residual eye size and %B across notothenioid species. We additionally repeated these PGLS regressions and all other downstream comparative phylogenetic analyses representing eye size using residuals from the regression of eye diameter on SL. Across all tests of correlated evolution, *p* values were adjusted using the false discovery rate corrections method of Benjamini and Hochberg (Benjamini and Hochberg 1995) using the p.adjust R function.

We visually assessed the relationships among eye size, depth occupancy, and buoyancy across the time-calibrated notothenioid phylogeny using a series of phenograms generated in the R package phytools ((Revell 2012)). These projections of a phylogeny in trait space have branch lengths corresponding to evolutionary time and the vertical positions of the tips corresponding to measured trait values. We used the contMap function of phytools to map evolution of mean depth of occurrence onto the projected phylogeny in order to simultaneously visualize patterns of evolution in eye size and depth occupancy along the notothenioid phylogeny. This procedure was repeated with %B mapped onto a phylogeny projected into trait space defined by eye size in order to compare evolutionary patterns of eye size and buoyancy.

To investigate patterns of morphospace occupancy over the course of the notothenioid radiation we next generated phylomorphospaces (Sidlauskas 2008) and also calculated trait subclade disparity through time (Harmon et al. 2003). First, we used the phylomorphospace function in the R package phytools ((Revell 2012)) to project the notothenioid phylogeny onto a biplot of mean depth of occurrence versus eye size. Values of internal nodes were estimated using a maximum likelihood-based approach implemented using the fastAnc function of phytools. This visualization enabled us to assess the directionality and magnitude of changes in both eye size and depth along each branch of the notothenioid phylogeny while also allowing us to evaluate the extent to which morphological similarity is predicted by phylogenetic relatedness. We additionally used fastAnc to map evolution of %B onto the phylomorphospace in order to facilitate visualization of evolutionary changes in buoyancy in relation to evolutionary changes in depth and in eye size.

We calculated relative subclade disparity through time (DTT; (Harmon et al. 2003)) to evaluate patterns of disparity over the course of the notothenioid radiation using the R package geiger2 (Pennell et al. 2014) for the traits of eye size, mean depth of occurrence, and %B. We assessed whether patterns of notothenioid trait disparity significantly departed from expectations under a null model of Brownian motion generated from 10,000 simulations of trait evolution on the notothenioid phylogeny. Divergence of the empirical DTT curve from the Brownian motion simulations was assessed using the morphological disparity index (MDI), with negative MDI values indicating that trait disparity is partitioned mostly among subclades, while positive MDI values suggest that trait disparity is partitioned within subclades. Following Harmon et al. (Harmon et al. 2003), we restricted our calculation of MDI values to only the first 80% of the notothenioid time tree. We further complemented our analysis of disparity with a quantification of the rate of eye size evolution using a Bayesian analyses of macroevolutionary mixtures (BAMM)(Rabosky 2014), with prior parameters set using the R package BAMMtools (Rabosky et al. 2014). This analysis allowed us to assess evidence for an elevated rate of eye size evolution during the initial radiation as would be predicted by classical adaptive radiation theory (Simpson 1953; Gavrilets and Losos 2009) versus evidence of more recent eye size diversification following more recent ecological opportunities generated by community recovery from glacial scouring events (Dornburg et al. 2017; Parker et al. 2022). We ran two independent analyses for 50 million generations, sampling every 1,000 generations. Convergence between runs was assessed through visual inspection of the log-likelihoods and effective sampling of the target posterior distribution of parameter values was assessed through quantification of ESS values (ESS >200). We assessed the sensitivity of our diversification rate analysis to the prior in BAMM (Moore et al. 2016; Rabosky et al. 2017) with replicate analyses favoring 0, 1, and 2 rate shifts.

In order to evaluate the extent to which variation in our focal ecomorphological traits can be explained by phylogeny, we additionally calculated phylogenetic signal in eye size, mean depth of occurrence, and mean %B. Specifically, we used the phylosig function implemented in the R package phytools ((Revell 2012)) to calculate Pagel’s λ (Pagel 1999) and Blomberg et al.’s *K* (Blomberg et al. 2003) for each trait based on the time-calibrated notothenioid phylogeny derived from Near et al. (Near et al. 2018). For Pagel’s λ, a value of zero indicates no phylogenetic signal is present in the data, and a value of one indicates that phylogenetic signal is consistent with expectations of a Brownian Motion model of trait evolution. The statistical significance of Pagel’s λ values was determined using a likelihood ratio test comparing the empirical value of Pagel’s λ to the null expectation that λ = 0. For Blomberg et al.’s *K*, values less than one suggest lower phylogenetic signal than expected under a BM model, while values greater than one suggest higher than expected phylogenetic signal. The statistical significance of Blomberg et al.’s *K* was determined by evaluating the degree to which empirical calculations of *K* deviated from a null distribution of expected *K* values from 10,000 permutations of trait evolution on the notothenioid tree.

## Results

### The evolution of cryonotothenioid opsin tuning sites

We identified opsin sequences from 25 notothenioids including 23 cryonotothenioids (Supplemental Tables S2 and S3) and confirmed their classification as Rh1, Rh2, SWS2, SWS1, or LWS through phylogenetic analyses (Figure 1). Our results reveal a complex history of tuning site substitutions that provide evidence for conserved changes that predate the onset of the cryonotothenioid adaptive radiation, changes that likely occurred coincident with the initial adaptive radiation, and numerous independent shifts towards lower frequency light wavelengths within cryonotothenioids. By comparing cryonotothenioid opsin sequences to other notothenioids along with perciforme and teleost outgroups, we identified a number of changes in icefish opsins that may influence their maximum wavelength of absorbed light λ_max_ (Figure 1, Supplemental Figures S1-S5 and Table S4).

**Figure 1.**
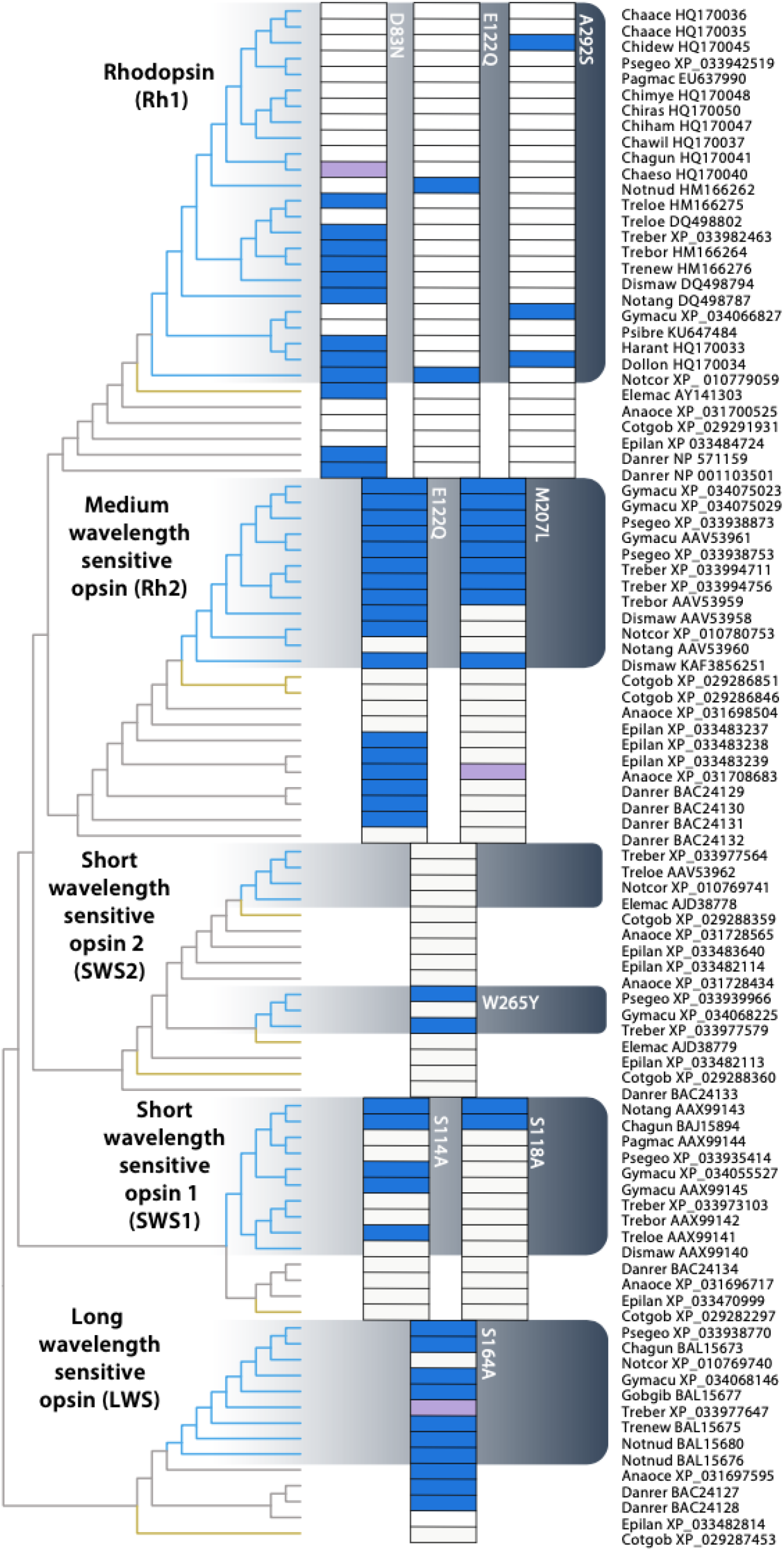
The evolution of opsin tuning site replacements in cryonotothenioids. Maximum likelihood topology of opsin sequences (left) with tuning site replacements indicated in the shaded grids. Clades identity is indicated on the phylogeny and cryonotothenoids are indicated by the gray gradient box and blue shading on the phylogeny. Purple boxes indicate alternate substitutions (see supplemental materials). Other nototheniods are indicated on the phylogeny in yellow, other teleost outgroups in gray. Text near the shaded boxes indicates AA replacement (*i.e*., W265Y). Names on the right provide sequence accession numbers and genus/species codes. Genus, species and common names are provided in Supplemental Table S2.

The four tuning sites of teleost Rh1 that, when mutated, can significantly shift the λ_max_ are D83 E122, F261 and A292 (Bowmaker 2008; Yokoyama 2008; Lin et al. 2017), and we observed multiple instances of D83N (−3 nm), and sporadic occurrences of E122Q (−13 nm) and A292S (−2 nm), as well as combined D83N and A292S substitutions (−14-17 nm) in cryonotothenioid Rh1 sequences (Figure 1, Supplemental Figure S1 and Table S4). We identified tuning site changes, E122Q and M207L, in the majority of cryonotothenioid Rh2 sequences, with some lineage specific losses in *Notothenia* and *Dissostichus* (Figure 1, Supplemental Figure S2 and Table S4). These substitutions could decrease λ_max_ of Rh2 by up to 17 and 10 nm (E122Q and M207L, respectively) (Yokoyama et al. 1999; Takenaka and Yokoyama 2007; Yokoyama and Jia 2020). In contrast to this signature of amino acid replacements during the early radiation of the clade in Rh2, the W265Y replacement in one copy of SWS2 revealed a convergence in a nearly 30 nm shift towards smaller light wavelengths (Yokoyama et al. 2007) that occurred independently in two distantly related species (*Pseudochaenichthys georgianus* and *Trematomus bernacchii* (Figure 1, Supplemental Figure S3 and Table S4). Similar independent convergences also occurred in SWS1 replacements S114A and S118A (Figure 1, Supplemental Figure S4 and Table S4). Finally, we identified two distinct changes in a single tuning site: S164A and S164P cryonotothenioid LWS sequences, the latter representing a unique substitution in *Trematomus bernacchi* (Figure 1, Supplemental Figure S5, and Supplemental Table S4). Collectively, almost all of these changes in cryonotothenioid opsins are predicted to result in decreases in λ_max_ with shifts towards shorter wavelengths (Supplemental Table S4).

### Ecological and morphological evolution

Analysis of depth data for cryonotothenioids reveal numerous instances of taxa transitioning to depths well beyond the photic zone (Figure 2). Simultaneous visualization of variation in mean depth occupancy and residual eye size corrected for body size across the cryonotothenioid phylogeny reveals that both the deepest- and shallowest-dwelling notothenioid species occupy overlapping extremes of variation in eye size (Figure 2). For example, *Pleurogramma* is often encountered deeper than 500m and possesses large eyes relative to body size, while the similarly depth distributed *Pogonophryne mantella* exhibits small eye size relative to body size (Figure 2). Correspondingly, results of our PGLS regression analysis revealed no significant evidence that eye size is correlated with mean depth of occurrence, minimum reported depth, or maximum reported depth across notothenioids (*p* > 0.05 for all regressions; Table 1). Contrasting these shifts in eye size and depth occupancy with tuning site replacements further reveals lack of a consistent ecomorphological signal. For example, *Trematomus loennbergii* and *Champsocephalus gunnari* have converged on the SWS1 replacement S114A (Figure 1). However, *Trematomus loennbergii* exhibits a substantial enlargement of eyes relative to body size and occupies depths of over 600m (Figure 2), while the distantly related *Champsocephalus gunnari* commonly occupies depths less than 300m and exhibits no dramatic eye size enlargement. Similarly, *Dolloidraco, Chionobathyscus*, and *Gymnodraco* all converge on an A292S replacement in Rh1 following transitions to deeper water (Figure 1 & 2). *Dolloidraco* and *Chionobathyscus* exhibit no substantial eye enlargement or reduction, while *Gymnodraco* ranks among the cryonotothenioids with the smallest eyes relative to body size (Figure 2).

**Figure 2.**
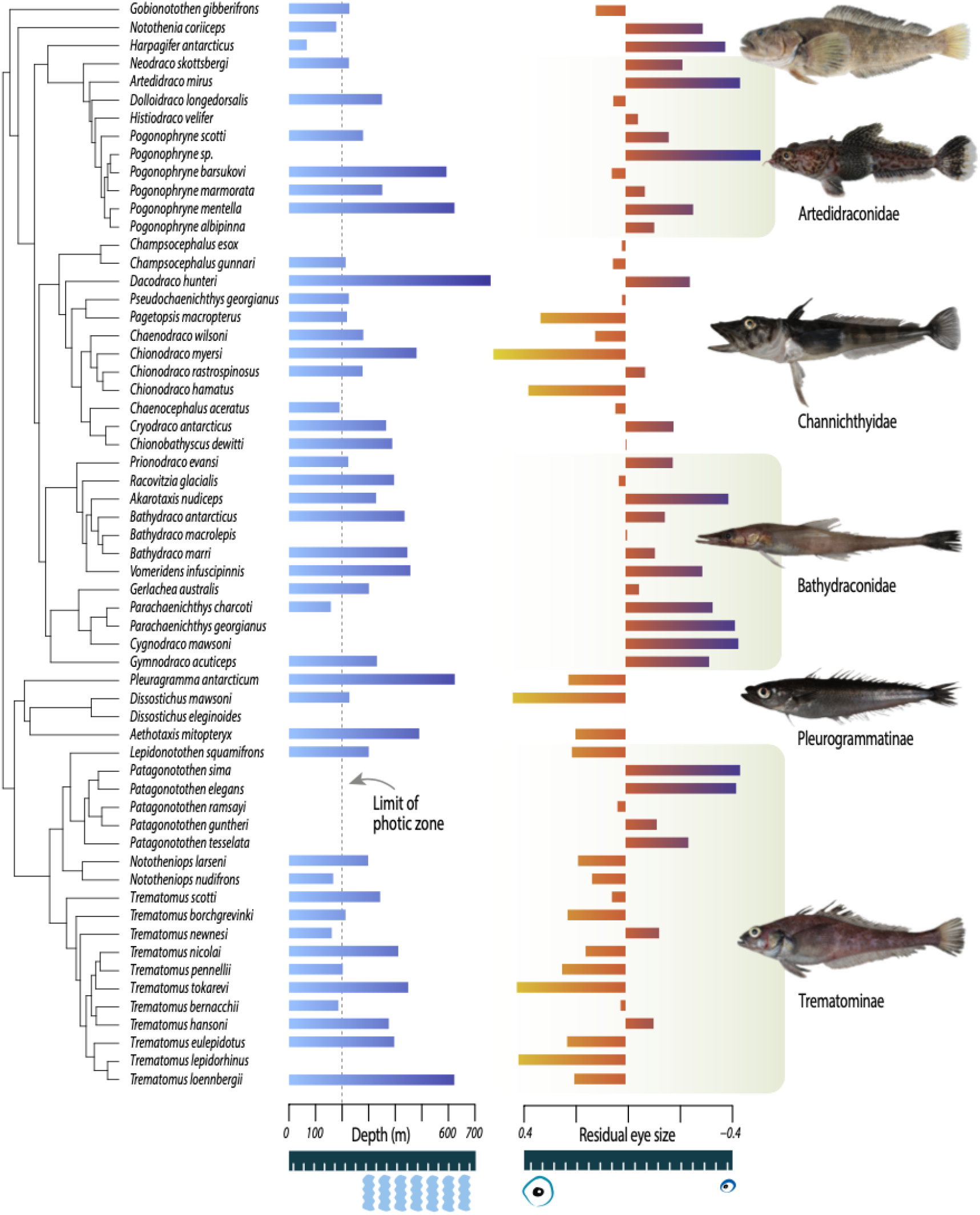
Visualization of variation in mean depth and eye size across the notothenioid phylogeny. Shown on the left panel is a time-calibrated tree derived from Near et al. (2018) depicting phylogenetic relationships among notothenioid species sampled in our morphological dataset. The middle panel depicts average depth per species, with darker shadings corresponding to deeper depths. The right panel depicts a barplot of eye size (represented as residuals from the regression of eye diameter on head shape) measured for our focal notothenioid species with warm colors representing larger eyes relative to head size. Fish images: EP.

**Table 1.** Results from PGLS regressions of residual eye size on depth and buoyancy for all notothenioid species.

| Predictor Variable | Parameter estimate<br>($\beta$) ± SE | t-value | p-value<sup>1</sup> |
| --- | --- | --- | --- |
| Mean depth of occurrence | 0.00016 ± 0.0012 | 0.13 | 0.90 |
| Minimum reported depth | -0.00080 ± 0.00088 | -0.91 | 0.90 |
| Maximum reported depth | -0.000078 ± 0.00035 | -0.22 | 0.90 |
| Mean % Buoyancy | -0.097 ± 0.18 | -0.53 | 0.90 |

1. P-values have been adjusted to correct for multiple comparisons using the false discovery rate correction method of Benjamini and Hochberg (1995).

We further find no significant evidence that shifts in eye size are associated with predictable shifts in %B (*p* > 0.05; Table 1). For instance, among notothenioid species with the largest relative eye sizes, we identify examples of both neutrally-buoyant pelagic species (e.g. *Pleuragramma antarcticum*) and comparatively heavy benthic species (e.g. *Nototheniops nudifrons*) (Figure 3A). Contrasting patterns of eye size evolution relative to either %B (Figure 3A) or depth (Figure 3B) reveals numerous examples of convergence in these habitat traits with divergences in eye size. These contrasts are most pronounced in channichthyids and trematomines (Figure 3 and Supplemental Figure S6), suggesting an asymmetry in the evolution of eye sizes between clades. In total, these results reveal extreme instances of convergence in eye size and depth occupancy across the evolutionary history of cryonothenioids (Figure 3).

**Figure 3.**
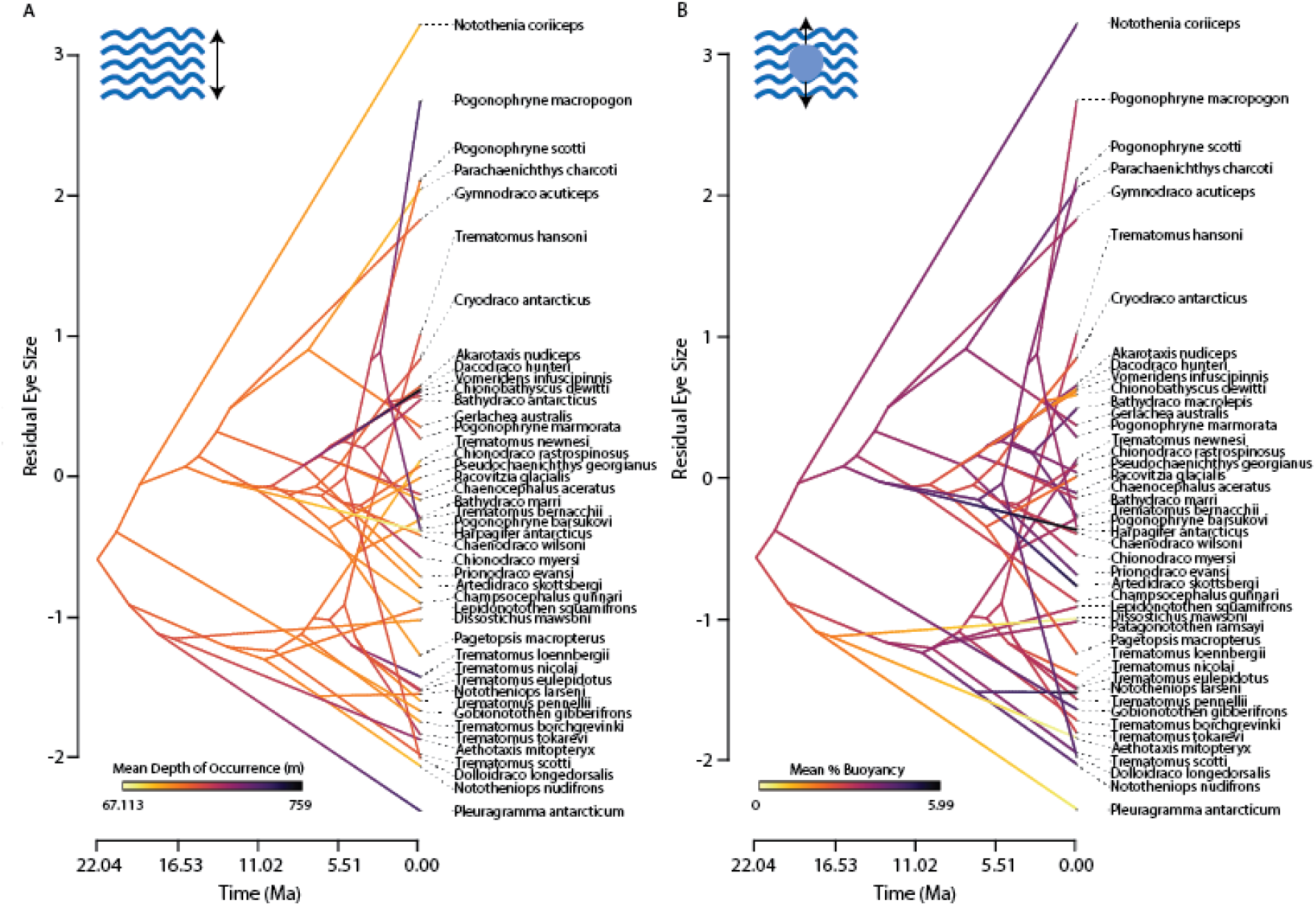
Projection of notothenioid phylogeny in space defined by time and eye size. Time (in millions of years) is on the X axis and eye size variability is on the Y axis. Placement of tree tips along the Y axis corresponds to eye size (represented using the residuals from regression of eye diameter on head length) for each notothenioid species. Ancestral state reconstructions of mean depth of occurrence (panel A) and of mean %B (panel B) have been mapped onto the notothenioid phylogeny to facilitate simultaneous visualization of variation in eye size and variation in ecology.

The decoupling of evolution in eye size, depth, and buoyancy is further underscored by analyses of relative subclade disparity in these traits through time (Figure 4). Both eye size and mean depth of occurrence depict a signature of higher within than between clade disparity (Figure 4A-B), corresponding to divergences between closely related species and convergences in eye size and depth niche between distantly related species. In contrast, %B generally follows the mean expectations of a brownian motion model, with no elevated within clade diversification (Figure 4C). This lack of within clade disparity in %B corresponds to strong phylogenetic signal (Table 2; Supplemental Figure S7), providing further support for similar %B among closely related species. Although there is moderate support for similar conservation between closely related taxa in eye size based on Pagel’s lambda statistic, this result is not significant based on Blomberg’s K (Table 2). Moreover, a Bayesian quantification of the rate of eye size diversification reveals a rapid and recent acceleration of eye size diversification through time (Figure 5). This sudden increase in diversification is driven largely by divergences in eye size between closely related taxa and occurs following the end of the mid-Pliocene warming, nearly 20 million years after the initial onset of the notothenioid adaptive radiation (Figure 5).

**Figure 4.**
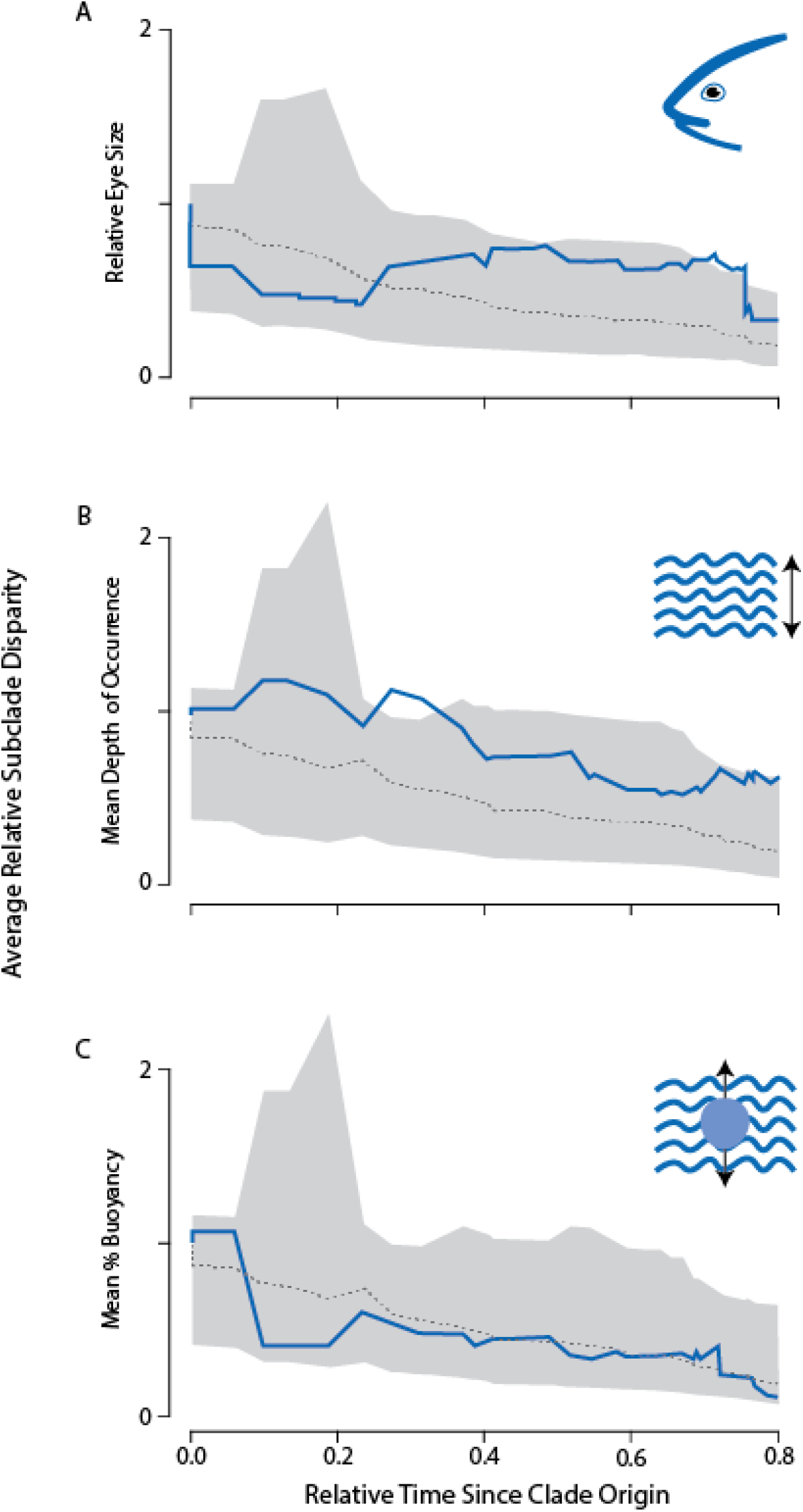
Disparity through time (DTT; Harmon et al. 2003) in eye size and ecology over the course of the notothenioid radiation. Panel A depicts patterns of disparity in residual eye size corrected for head length, panel B reflects disparity in mean depth of occurrence, and panel C depicts disparity in mean % buoyancy. In all plots, the X axis reflects relative time since clade origin (0.0). The Y axis corresponds to average relative subclade disparity in eye size. The solid blue line depicts the empirical estimation of eye size disparity, while the dotted gray line depicts the median trait disparity calculated from 10,000 Brownian motion simulations of trait evolution on the notothenioid phylogeny. The shaded gray region represents the 95% confidence interval (CI) of the Brownian motion simulations.

**Figure 5.**
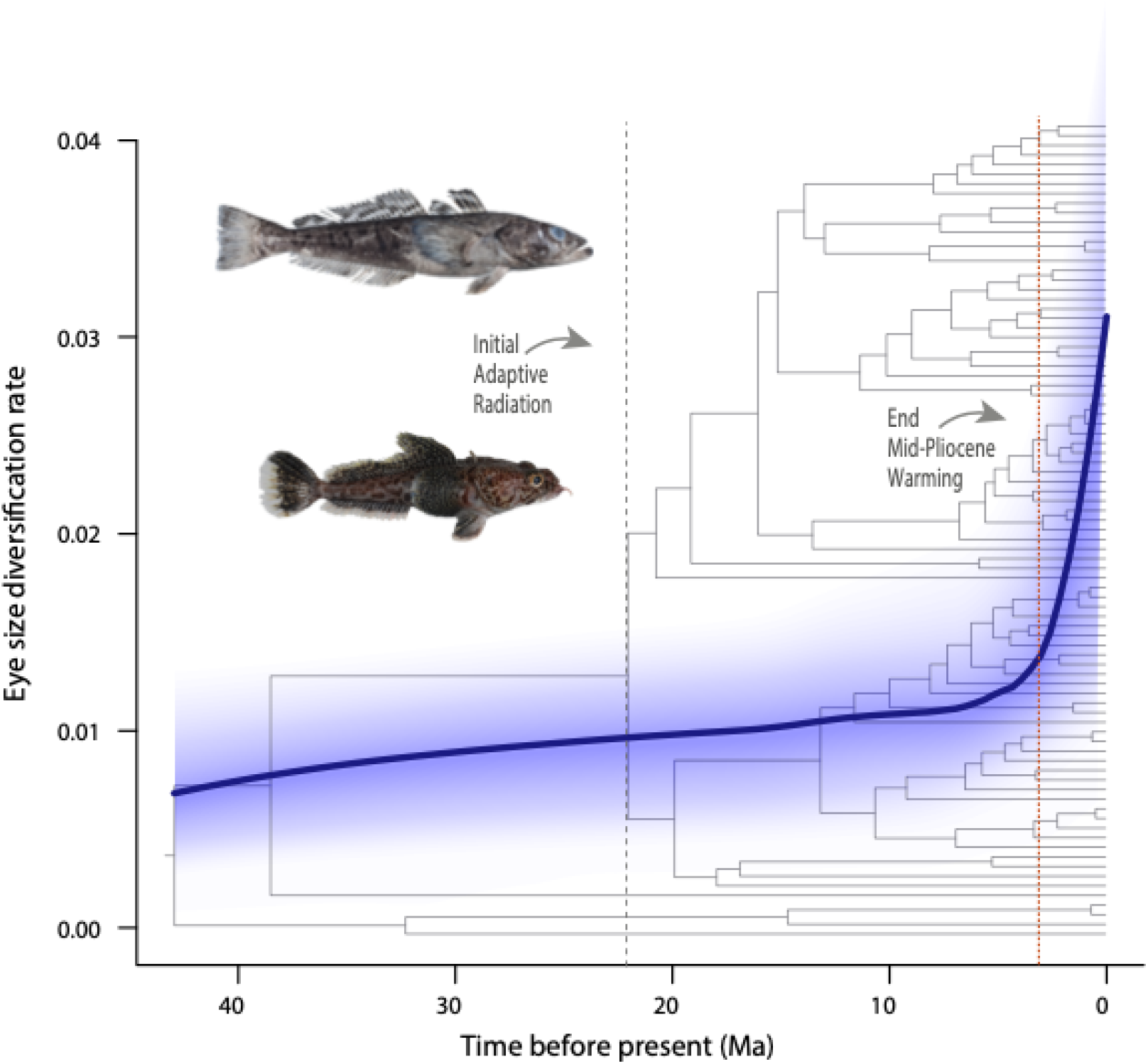
Punctuated elevation in the diversification of eye size well after the onset of the cryonotothenioid adaptive radiation. The mean rate of eye size diversification (solid line) and confidence interval (blue shading) overlaid on the time calibrated phylogeny of notothenioids reveals a signature of a punctuated increase in eye size diversification coincident with the end of the mid-pliocene warming (orange line) that occurred nearly 20 Ma after the initial antarctic radiation (gray line; (Near et al. 2012; Daane et al. 2020)). Photos: EP.

**Table 2.** Results of tests of phylogenetic signal in ecomorphological traits.

| Traits | Pagel's lambda |  | Blomberg et al.'s K |  |
| --- | --- | --- | --- | --- |
|  | Estimate | *p*-value | Estimate | *p*-value |
| Residual Eye Size | 0.36 | **0.0016** | 0.35 | 0.27 |
| Mean Depth of Occurrence | 7.3E-05 | 1 | 0.43 | 0.28 |
| Mean % Buoyancy | 1.04 | **5.60E-06** | 0.88 | **1.0E-04** |

## Discussion

For a closely related group of species, evolutionary radiation across the water column may be expected to require a correlation between the diversification of depth niche, eye size, and the molecular basis of light detection (Huber et al. 1997). However, our results strongly suggest such an expectation does not characterize the Antarctic notothenioid adaptive radiation. Instead, we reveal a complex history of trait convergences and divergences that collectively suggest evolutionary changes in opsin tuning sites, eye size, depth niche, and buoyancy to be largely decoupled in this radiation. We find a consistent signature of shifts in tuning sites reflecting shifts towards lower wavelengths of light that represent repeated instances of independent tuning site changes among distantly related species that are not necessarily associated with convergences in depth or eye size. Our analysis further reveals that convergences in depth result in no clear association with eye size or buoyancy. Lineages occupying similar depths give rise to extremes in eye size and often depict different levels of buoyancy. Eye size divergence is often high among closely related taxa, and our results further reveal that the rate of eye size diversification in notothenioids accelerated nearly 20 million years after the initial radiation, likely in response to recovery from the end of the mid-Pliocene warming event. Collectively, our results reveal that the diversification of these key components of the visual system do not conform to expectations of the classic adaptive radiation model.

### The mosaic evolution of cryonotothenioid opsins

Closely related notothenioid species often diverge in their water column niches by hundreds of meters, subjecting them to new ecological opportunities and photic conditions (Parker et al. 2022). While it has been established that changes in opsin tuning sites can result in increases or decreases in the maximum wavelength of absorbed light (λ_max_) (Lin et al. 2017; Bowmaker 2008; Yokoyama 2008), changes in opsin sites are often not consistent relative to depth niche between cryonotothenioid species. Likewise, there is no evidence that changes in eye size correspond to consistent changes in tuning sites. Instead, cryonotothenioids have maintained some tuning sites that likely arose prior or coincident with their early radiation as well as with numerous substitutions that arose throughout the radiation in various lineages. For example, it is expected that the observed E122Q substitution within Rh2 of notothenioids would decrease λ_max_ by 13-17 nm (Takenaka and Yokoyama 2007). The E122Q substitution is found in nearly all cryonotothenioid Rh2 sequences we investigated (with the exception of a reversal in *Notothenia angustata*). This substitution is also found in a wide range of teleost lineages including other perciformes (Lin et al. 2017), indicating that the E122 in *Cottoperca* could reflect a reversal. The M207L mutation in Rh2 is also pervasive across most surveyed cryonotothenioids, and predicted to decrease λ_max_ by 6 nm (Yokoyama et al. 1999). As the M207L mutation has been found in only a few teleost lineages (e.g. *Fundulus, Xiphophorus* and *Poecilia*) (Lin et al. 2017) but not in other perciformes included in this study (Supplemental Figure S2), this suggests the M207L mutation may be a unique feature of cryonotothenioids.

In contrast to these widespread substitutions, we observe the combination of D83N with A292S in the Rh1 of *Chionobathyscus dewitti* and *Gymnodraco acuticeps* that likely reflect a λ_max_ decrease of ~13-17 nm (Yokoyama et al. 2008). The A292S change has previously been identified as an adaption for deeper water in cichlids (Sugawara et al. 2005), and the instance of this substitution in *Chionobathyscus dewitti* and *Gymnodraco acuticeps* likely reflects an evolutionary convergence in these deeper dwelling taxa. However, these mutations are not uniform in all deep dwelling taxa occurring at similar or deeper depths. We also observe an E122Q mutation in Rh1 in the distantly related *Nototheniops nudifrons* and *Notothenia coriiceps* that likely reflect a similar λ_max_ decrease of ~13 nm (Yokoyama et al. 2008). In a recent survey of ray-finned fish opsins, the E122Q mutation was found in *Notothenia coriiceps* (the only notothenioid in their report) and three other teleosts: Pacific bluefin tuna (*Thunnus orientalis*), yellowhead catfish (*Tachysurus fulvidraco*) and red piranha (*Pygocentrus nattereri*) (Lin et al. 2017). Similarly, the double mutation of D83N/A292S was only observed in a few other teleost lineages: red piranha (*Pygocentrus nattereri*), blind cave fish (*Astyanax mexicanus*), the eel species *Anguilla anguilla* and *A. japonica*, and ballan wrasse (*Labrus bergylta*) (Lin et al. 2017). The rarity of these substitutions across all ray-finned fish lineage compared with the frequency of our observations in cryonotothenioids suggests these to likely be more widespread across other cryonotothenioids and an exciting area of future functional study.

To date, no report has identified all opsins from an individual notothenioid species and little information has been reported about their tuning sites. While we confirm and expand the scope of changes previously identified in a limited number of notothenioids, such as those in SWS1 (S114A; S118A), Rh2 (E122Q; M207L) (Pointer et al. 2005), LWS (S164A) (Miyazaki and Iwami 2012), and Rh1 (E122Q)) (Lin et al. 2017), we also identified an undescribed notothenioid substitution that exceeds the spectral shift of any previously identified substitution. We find a W265Y substitution in SWS2 sequences in the distantly related *Pseudochaenichthys georgianus* and *Trematomus bernacchii* that is predicted to lead to a significant decrease in λ_max_ by 29 nm (Yokoyama et al. 2007). This substitution has been observed in only a few teleost lineages (*e.g*., killifish), where there are two copies of SWS2 where SWS2-A retains W265 and blue-sensitivity, and SWS2-B provides violet-sensitivity through a Y265 substitution (Yokoyama et al. 2007; Lin et al. 2017). Our analyses suggest that *Trematomus bernacchii* encodes SWS2-A and SWS2-B, and more sequencing is needed from other notothenioids to determine if all notothenioids encode both SWS2 opsins. If W265Y is indeed more pervasive across cryonotothenioids, this would suggest a substantial shift towards shorter blue/UV light wavelengths and further underscore a key result of our study: functional changes in cryonotothenioid opsins represent an evolutionary mosaic shifted towards shorter light wavelengths.

### Evolving to gaze into the abyss

Changes in water column usage are a major axis of diversification in ray-finned fishes (Ingram 2011; Iglesias et al. 2015; Tavera et al. 2018), and the stratification of cryonotothenioids across the water column of the Southern Ocean is a hallmark of their evolutionary success (Eastman and McCUNE 2000; Daane et al. 2019; Parker et al. 2022). However, we find no strong evidence for coordinated evolutionary changes between opsin tuning sites, eye size, depth niche, or buoyancy. Instead, we find repeated convergences in depth niches and eye size diversity between lineages, suggesting that these traits are highly labile. In particular, relative eye size changes appear most pronounced within Channichthyidae and Trematominae, two lineages previously identified as nested adaptive radiations (Near et al. 2012; Parker et al. 2022). In these clades, our analyses reveal closely related taxa to often sharply diverge in eye size, resulting in a high degree of eye size convergence between distantly related taxa. This pattern of recent divergences is also apparent in our quantification of the rate of eye size evolution, and likely reflects the recovery of lineages following ice scour events that decimated near-shore ecological communities (Thatje et al. 2005, 2008; Dornburg et al. 2017; Parker et al. 2022). As lineages colonize recovering communities, stratification of the vertical water column presents a mechanism to mitigate resource conflict and explore additional ecological opportunities (La Mesa et al. 2004a). The frequency of these transitions may derive from a predisposition to dim-light vision in cryonotothenioids. The depth ranges occupied by notothenioids tend to be deeper than most ‘near-shore’ fauna given the depression of continental shelf and polar latitudes experience extreme seasonal fluctuations in sunlight along with shifts in light attenuation due to ice coverage and phytoplankton biomass (Tilzer et al. 1994). Such conditions have likely primed notothenioids to repeatedly converge in depth occupancy patterns across hundreds of meters in short evolutionary timescales.

Larger eyes are associated with an increase in photoreceptors that can collect higher amounts of sensory information (Iglesias et al. 2018) While such increases could be expected to aid lineages in prey detection, we found that the evolution of eye size and depth niche are not correlated in notothenioids. We suggest that this lack of correlation between eye size and depth niche may reflect trade-offs in the neurological investment between chemo perception and visual acuity. Across vertebrates, eye size has repeatedly been shown to be linked with aspects of brain architecture, especially those associated with vision (Burton 2008; Corral-López et al. 2017; Howell et al. 2021). Many predatory teleosts that hunt in dim-light conditions have reduced eye sizes and correspondingly reduced optic tecta, instead relying on olfaction and concomitant increases in the size of the brain’s olfactory bulb for prey detection (Edmunds et al. 2016; Yamamoto 2017; Iglesias et al. 2018). In contrast, fishes that constitute the prey of such lineages foraging in open environments often rely on visual acuity to detect incoming motion and ambush predators (Guthrie 1990; Kotrschal et al. 2017). As some predatory notothenioids are themselves primary prey items for hunting marine mammals and birds (Eastman 1985,La Mesa et al. 2004b; Casaux et al. 2011), predation pressure could be a major force shaping the evolution of notothenioid eye size and associated investment in corresponding brain regions. Such a hypothesis would be in line with growing evidence that predation pressure can impact the evolution of the brain within (White and Brown 2015)and between species (Kotrschal et al. 2017). Future studies contrasting neural investment patterns with ecological data across the phylogeny of notothenioids are needed to assess the degree to which eye size and brain evolution have been impacted by predation pressure.

An additional key trait to the notothenioid radiation in the water column is their ability to modulate buoyancy. As all notothenioids lack a swim bladder, changes in skeletal ossification permit notothenioids to utilize deeper or shallower water column niches (Daane et al. 2019; Daane and William Detrich 2021). However, we find a surprising lack of correlation between buoyancy and water column niche. This lack of correlation is likely explained by evolutionary transitions along the benthic-pelagic axis of the water column. Benthic species will naturally be more negatively buoyant and co-occur with pelagic species across the depth range of notothenioids. Unfortunately, understanding the relationship between traits in this study and this aspect of the water column niche is complicated by uncertainty in classification of notothenioid species into benthic or pelagic categories. Some species, such as *Pleuragramma antarcticus*, *Dissostichus mawsoni*, *D. eleginoides*, and *Aethotaxis mitopteryx*, are unquestionably pelagic (Eastman and DeVries 1982; Eastman 2020) (Near et al. 2003, 2007), and others, such as species of *Harpagifer*, can almost certainly be considered benthic (Eastman 2020). In addition, classifying the majority of notothenioid species into water column niches is complicated by limited ecological data as well as the behavioral plasticity that has been recorded for notothenioids. For instance, there are several notothenioids that have been observed to opportunistically feed “outside” of their “expected” water column niche including benthic species opportunistically capturing pelagic prey (Daniels 1982). Continual ecological investigations focused on how notothenioids are utilizing their habitat are needed to place our analyses into a more refined ecological context, as it is currently not possible to accurately contrast patterns of eye size or opsin evolution relative to these aspects of water column niches. Fortunately, continual advances in imaging technology hold the promise of overcoming traditional logistical challenges of observing notothenioids in the Antarctic (Jones and Near 2012; La Mesa et al. 2021; Purser et al. 2022) and providing these critically needed insights into the ecology of this adaptive radiation in the near future.

### Conclusion

It has become clear over the last decade that the adaptive radiation of notothenioids is far more complex than previously hypothesized. Rather than merely representing a clade that radiated at the onset of polar conditions in the Antarctic over 30 million years ago (Eastman and McCUNE 2000; Rutschmann et al. 2011), this radiation is comprised of nested radiations that have arisen in response to recent warming events in the Plio/Pleistocene and ongoing cycles of ice scouring across the continental shelf (Near et al. 2012; Dornburg et al. 2017; Parker et al. 2022). Our results are consistent with this emerging view, demonstrating an acceleration of eye size evolution following the mid-Pliocene warming event, and high levels of ecological and morphological divergence between closely related species. We further reveal that the diversification of notothenioid opsins reflects a complex evolutionary mosaic that includes both molecular substitutions in tuning sites that occurred early in the group’s history as well as recent reversals and novel substitutions. Collectively, patterns of depth usage, tuning site substitutions, or eye sizes are heterogeneous even among close evolutionary relatives with convergences in taxa that are otherwise highly divergent in other aspects of life history. This decoupling of ecological, morphological, and genotypic traits raises the possibility that the relationship between ecological functions and visual acuity may represent a many-to-one mapping of forms to function, similar to that often observed in studies of feeding performance and morphology (Alfaro et al. 2005; Wainwright et al. 2005). As changes in eye size and visual acuity are often correlated with changes in neural investment (Iglesias et al. 2018; Howell et al. 2021), future comparative considerations of transitions in depth and the compositional diversity of notothenioid brains offers an exciting research frontier that can illuminate the pathways and trade-offs that underlie neural diversification in this adaptive radiation. Given the environmental threats facing the Southern Ocean (Patarnello et al. 2011; Vacchi et al. 2017), such studies are also of high conservation importance for understanding the impact of warming surface waters on lineages predicted to alter their depth niches in response.

## Author Contributions

EBY and AD designed the project. EBY identified, aligned and analyzed opsin sequences and tuning sites. AT assisted with identifying opsin sequences. AT, EP, and AD collected eye size measurements and conducted comparative analyses. EBY, AD, and EP wrote the first draft of the manuscript. CDJ led finfish trawl surveys, reviewed and edited the manuscript. All authors read and approved the final manuscript.

## Acknowledgments

We thank members of the Dornburg lab for support and feedback during early project stages.

## Data Accessibility Statement

All data generated or analyzed during this study are included in this published article (and its supplementary information files). Code, data, and trees to replicate analyses are available on Zenodo: 10.5281/zenodo.5106397

## Funding

NA

## Declarations

### Ethics approval and consent to participate

Not applicable

### Consent for publication

Not applicable

### Competing interests

The authors declare that they have no competing interests

## Notes

https://zenodo.org/record/7116758#.YzQqGOzMIeY

### Competing Interest Statement

The authors have declared no competing interest.

