## Supplemental materials for "Decoupled spectral tuning and eye size diversification patterns in an Antarctic adaptive radiation"

**Table S1. List of specimens used for morphological analyses.**

| Clade | Genus species | Museum Voucher[1] |
|---|---|---|
| Bathydraconidae | *Akarotaxis nudiceps* | YPM ICH 24241 |
| Bathydraconidae | *Akarotaxis nudiceps* | YPM ICH 20210 |
| Bathydraconidae | *Akarotaxis nudiceps* | YPM ICH 20037 |
| Bathydraconidae | *Bathydraco antarcticus* | NMNZ P043669 |
| Bathydraconidae | *Bathydraco antarcticus* | NMNZ P043670 |
| Bathydraconidae | *Bathydraco macrolepis* | NMNZ P042270 |
| Bathydraconidae | *Bathydraco macrolepis* | NMNZ P043387 |
| Bathydraconidae | *Bathydraco marri* | NMNZ P040098 |
| Bathydraconidae | *Bathydraco marri* | YPM ICH 22590 |
| Bathydraconidae | *Bathydraco marri* | YPM ICH 22477 |
| Bathydraconidae | *Cygnodraco mawsoni* | NMNZ P040018 |
| Bathydraconidae | *Gerlachea australis* | YPM ICH 16476 |
| Bathydraconidae | *Gerlachea australis* | YPM ICH 16642 |
| Bathydraconidae | *Gerlachea australis* | NMNZ P043338 |
| Bathydraconidae | *Gymnodraco acuticeps* | YPM ICH 16635 |
| Bathydraconidae | *Gymnodraco acuticeps* | YPM ICH 22376 |
| Bathydraconidae | *Gymnodraco acuticeps* | NMNZ P043639 |
| Bathydraconidae | *Parachaenichthys charcoti* | YPM ICH 22649 |
| Bathydraconidae | *Parachaenichthys charcoti* | YPM ICH 22649 |
| Bathydraconidae | *Parachaenichthys charcoti* | YPM ICH 22649 |
| Bathydraconidae | *Parachaenichthys georgianus* | YPM ICH 21533 |
| Bathydraconidae | *Parachaenichthys georgianus* | YPM ICH 21533 |
| Bathydraconidae | *Parachaenichthys georgianus* | YPM ICH 21533 |
| Bathydraconidae | *Prionodraco evansii* | YPM ICH 22536 |
| Bathydraconidae | *Prionodraco evansii* | YPM ICH 22689 |
| Bathydraconidae | *Prionodraco evansii* | YPM ICH 24283 |
| Bathydraconidae | *Racovitzia glacialis* | YPM ICH 16660 |
| Bathydraconidae | *Racovitzia glacialis* | YPM ICH 20044 |
| Bathydraconidae | *Racovitzia glacialis* | YPM ICH 22370 |
| Bathydraconidae | *Vomeridens infuscipinnis* | YPM ICH 22391 |
| Bathydraconidae | *Vomeridens infuscipinnis* | YPM ICH 22391 |
| Bathydraconidae | *Vomeridens infuscipinnis* | YPM ICH 22391 |
| Channichthyidae | *Chaenocephalus aceratus* | YPM ICH 24284 |
| Channichthyidae | *Chaenocephalus aceratus* | YPM ICH 24284 |
| Channichthyidae | *Chaenocephalus aceratus* | YPM ICH 24284 |
| Channichthyidae | *Chaenodraco wilsoni* | YPM ICH 24286 |
| Channichthyidae | *Chaenodraco wilsoni* | YPM ICH 21474 |
| Channichthyidae | *Chaenodraco wilsoni* | YPM ICH 16641 |
| Channichthyidae | *Champsocephalus esox* | no tag |

[1] Institutional museum collection acronyms follow Sabaj (2016).

| Clade | Genus species | Museum Voucher[1] |
|---|---|---|
| Channichthyidae | *Champsocephalus gunnari* | YPM ICH 21535 |
| Channichthyidae | *Champsocephalus gunnari* | YPM ICH 21536 |
| Channichthyidae | *Champsocephalus gunnari* | YPM ICH 20242 |
| Channichthyidae | *Chionobathyscus dewitti* | YPM ICH 20010 |
| Channichthyidae | *Chionobathyscus dewitti* | YPM ICH 20011 |
| Channichthyidae | *Chionobathyscus dewitti* | YPM ICH 24123 |
| Channichthyidae | *Chionodraco hamatus* | no tag |
| Channichthyidae | *Chionodraco hamatus* | no tag |
| Channichthyidae | *Chionodraco myersi* | YPM ICH 22339 |
| Channichthyidae | *Chionodraco myersi* | YPM ICH 16533 |
| Channichthyidae | *Chionodraco rastrospinosus* | YPM ICH 22341 |
| Channichthyidae | *Chionodraco rastrospinosus* | YPM ICH 22341 |
| Channichthyidae | *Chionodraco rastrospinosus* | YPM ICH 20052 |
| Channichthyidae | *Cryodraco antarcticus* | YPM ICH 22605 |
| Channichthyidae | *Cryodraco antarcticus* | YPM ICH 16521 |
| Channichthyidae | *Cryodraco antarcticus* | YPM ICH 22492 |
| Channichthyidae | *Dacodraco hunteri* | YPM ICH 22338 |
| Channichthyidae | *Dacodraco hunteri* | YPM ICH 16569 |
| Channichthyidae | *Pagetopsis macropterus* | YPM ICH 22335 |
| Channichthyidae | *Pagetopsis macropterus* | YPM ICH 16510 |
| Channichthyidae | *Pagetopsis macropterus* | YPM ICH 20477 |
| Channichthyidae | *Pseudochaenichthys georgianus* | YPM ICH 22601 |
| Channichthyidae | *Pseudochaenichthys georgianus* | YPM ICH 22524 |
| Channichthyidae | *Pseudochaenichthys georgianus* | YPM ICH 16637 |
| Harpagiferidae | *Artedidraco mirus* | YPM ICH 23278 |
| Harpagiferidae | *Artedidraco mirus* | YPM ICH 23278 |
| Harpagiferidae | *Artedidraco skottsbergi* | NMNZ P043365 |
| Harpagiferidae | *Artedidraco skottsbergi* | NMNZ P040034 |
| Harpagiferidae | *Artedidraco skottsbergi* | YPM ICH 22372 |
| Harpagiferidae | *Dolloidraco longedorsalis* | YPM ICH 20033 |
| Harpagiferidae | *Dolloidraco longedorsalis* | YPM ICH 20038 |
| Harpagiferidae | *Dolloidraco longedorsalis* | YPM ICH 20030 |
| Harpagiferidae | *Harpagifer antarcticus* | YPM ICH 20810 |
| Harpagiferidae | *Harpagifer antarcticus* | YPM ICH 16630 |
| Harpagiferidae | *Harpagifer antarcticus* | YPM ICH 16630 |
| Harpagiferidae | *Histiodraco velifer* | NMNZ P043412 |
| Harpagiferidae | *Pogonophryne albipinna* | NMNZ P042633 |
| Harpagiferidae | *Pogonophryne barsukovi* | NMNZ P043539 |
| Harpagiferidae | *Pogonophryne barsukovi* | YPM ICH 16518 |
| Harpagiferidae | *Pogonophryne barsukovi* | YPM ICH 16518 |
| Harpagiferidae | *Pogonophryne marmorata* | YPM ICH 16550 |
| Harpagiferidae | *Pogonophryne marmorata* | YPM ICH 20496 |

| Clade | Genus species | Museum Voucher[1] |
|---|---|---|
| Harpagiferidae | *Pogonophryne marmorata* | YPM ICH 16550 |
| Harpagiferidae | *Pogonophryne mentella* | NMNZ P042618 |
| Harpagiferidae | *Pogonophryne mentella* | YPM ICH 20495 |
| Harpagiferidae | *Pogonophryne mentella* | YPM ICH 22565 |
| Harpagiferidae | *Pogonophryne scotti* | NMNZ P045673 |
| Harpagiferidae | *Pogonophryne scotti* | NMNZ P043419 |
| Harpagiferidae | *Pogonophryne scotti* | NMNZ P043370 |
| Harpagiferidae | *Pogonophryne* sp. nov. | NMNZ P042640 |
| Nototheniidae | *Aethotaxis mitopteryx* | NMNZ P043384 |
| Nototheniidae | *Aethotaxis mitopteryx* | NMNZ P043377 |
| Nototheniidae | *Aethotaxis mitopteryx* | YPM ICH 22552 |
| Nototheniidae | *Dissostichus eleginoides* | NMNZ P037795 |
| Nototheniidae | *Dissostichus eleginoides* | YPM ICH 17045 |
| Nototheniidae | *Dissostichus mawsoni* | NMNZ P042631 |
| Nototheniidae | *Dissostichus mawsoni* | YPM ICH 20787 |
| Nototheniidae | *Dissostichus mawsoni* | YPM ICH 20787 |
| Nototheniidae | *Gobionotothen gibberifrons* | YPM ICH 16629 |
| Nototheniidae | *Gobionotothen gibberifrons* | YPM ICH 20779 |
| Nototheniidae | *Gobionotothen gibberifrons* | YPM ICH 20724 |
| Nototheniidae | *Lepidonotothen squamifrons* | YPM ICH 22548 |
| Nototheniidae | *Lepidonotothen squamifrons* | YPM ICH 20490 |
| Nototheniidae | *Lepidonotothen squamifrons* | YPM ICH 20490 |
| Nototheniidae | *Notothenia coriiceps* | NMNZ P040097 |
| Nototheniidae | *Notothenia coriiceps* | NMNZ P040096 |
| Nototheniidae | *Notothenia coriiceps* | NMNZ P040096 |
| Nototheniidae | *Nototheniops larseni* | YPM ICH 24132 |
| Nototheniidae | *Nototheniops larseni* | YPM ICH 24132 |
| Nototheniidae | *Nototheniops larseni* | YPM ICH 24132 |
| Nototheniidae | *Nototheniops nudifrons* | YPM ICH 21496 |
| Nototheniidae | *Nototheniops nudifrons* | YPM ICH 21496 |
| Nototheniidae | *Nototheniops nudifrons* | YPM ICH 21496 |
| Nototheniidae | *Patagonotothen elegans* | YPM ICH 21588 |
| Nototheniidae | *Patagonotothen elegans* | YPM ICH 20076 |
| Nototheniidae | *Patagonotothen elegans* | YPM ICH 20076 |
| Nototheniidae | *Patagonotothen guntheri* | YPM ICH 20072 |
| Nototheniidae | *Patagonotothen guntheri* | YPM ICH 20077 |
| Nototheniidae | *Patagonotothen guntheri* | YPM ICH 20077 |
| Nototheniidae | *Patagonotothen ramsayi* | YPM ICH 20124 |
| Nototheniidae | *Patagonotothen ramsayi* | YPM ICH 20124 |
| Nototheniidae | *Patagonotothen ramsayi* | YPM ICH 20124 |
| Nototheniidae | *Patagonotothen sima* | YPM ICH 20125 |
| Nototheniidae | *Patagonotothen sima* | YPM ICH 20125 |

| Clade | Genus species | Museum Voucher[1] |
|---|---|---|
| Nototheniidae | *Patagonotothen sima* | YPM ICH 20125 |
| Nototheniidae | *Patagonotothen tessellata* | YPM ICH 20071 |
| Nototheniidae | *Patagonotothen tessellata* | YPM ICH 20071 |
| Nototheniidae | *Patagonotothen tessellata* | YPM ICH 20071 |
| Nototheniidae | *Pleuragramma antarcticum* | NMNZ P043380 |
| Nototheniidae | *Pleuragramma antarcticum* | YPM ICH 16405 |
| Nototheniidae | *Pleuragramma antarcticum* | YPM ICH 16405 |
| Nototheniidae | *Trematomus bernacchii* | YPM ICH 22595 |
| Nototheniidae | *Trematomus bernacchii* | YPM ICH 22400 |
| Nototheniidae | *Trematomus bernacchii* | YPM ICH 16652 |
| Nototheniidae | *Trematomus borchgrevinki* | TA537PAB01 |
| Nototheniidae | *Trematomus eulepidotus* | YPM ICH 16496 |
| Nototheniidae | *Trematomus eulepidotus* | YPM ICH 16496 |
| Nototheniidae | *Trematomus eulepidotus* | YPM ICH 16496 |
| Nototheniidae | *Trematomus hansoni* | YPM ICH 20463 |
| Nototheniidae | *Trematomus hansoni* | YPM ICH 22685 |
| Nototheniidae | *Trematomus hansoni* | YPM ICH 22685 |
| Nototheniidae | *Trematomus lepidorhinus* | NMNZ P057959 |
| Nototheniidae | *Trematomus lepidorhinus* | NMNZ P043658 |
| Nototheniidae | *Trematomus lepidorhinus* | NMNZ P043594 |
| Nototheniidae | *Trematomus loennbergii* | YPM ICH 20018 |
| Nototheniidae | *Trematomus loennbergii* | YPM ICH 20019 |
| Nototheniidae | *Trematomus loennbergii* | YPM ICH 20020 |
| Nototheniidae | *Trematomus newnesi* | YPM ICH 20492 |
| Nototheniidae | *Trematomus newnesi* | YPM ICH 20492 |
| Nototheniidae | *Trematomus newnesi* | YPM ICH 20492 |
| Nototheniidae | *Trematomus nicolai* | YPM ICH 18359 |
| Nototheniidae | *Trematomus nicolai* | YPM ICH 18359 |
| Nototheniidae | *Trematomus nicolai* | YPM ICH 18359 |
| Nototheniidae | *Trematomus pennellii* | YPM ICH 24242 |
| Nototheniidae | *Trematomus pennellii* | YPM ICH 24242 |
| Nototheniidae | *Trematomus pennellii* | YPM ICH 16507 |
| Nototheniidae | *Trematomus scotti* | YPM ICH 24279 |
| Nototheniidae | *Trematomus scotti* | YPM ICH 24279 |
| Nototheniidae | *Trematomus scotti* | YPM ICH 24279 |
| Nototheniidae | *Trematomus tokarevi* | YPM ICH 22481 |
| Nototheniidae | *Trematomus tokarevi* | YPM ICH 20027 |
| Nototheniidae | *Trematomus tokarevi* | YPM ICH 22408 |

#### Table S2. Genus, species, abbreviations, and common names in Figures 1 and S1-S5

| Genus species | Abbreviation | Group[2] | Common name[3] |
|---|---|---|---|
| *Anarrhichthys ocellatus* | *Anaoce* | P | wolf eel |
| *Chaenocephalus aceratus* | *Chaace* | C | blackfin icefish |
| *Chaenodraco wilsoni* | *Chawil* | C | spiny icefish |
| *Champsocephalus esox* | *Chaeso* | C | pike icefish |
| *Champsocephalus gunnari* | *Chagun* | C | mackerel icefish |
| *Chionobathyscus dewitti* | *Chidew* | C | NA |
| *Chionodraco hamatus* | *Chiham* | C | NA |
| *Chionodraco myersi* | *Chimye* | C | Myers' icefish |
| *Chionodraco rastrospinosus* | *Chiras* | C | ocellated icefish |
| *Cottoperca gobio* | *Cotgob* | N | channel bull blenny |
| *Danio rerio* | *Danrer* | T | zebrafish |
| *Dissostichus mawsoni* | *Dismaw* | C | Antarctic toothfish |
| *Dolloidraco longedorsalis* | *Dollon* | C | NA |
| *Eleginops maclovinus* | *Elemac* | N | Patagonian blennie |
| *Epinephelus lanceolatus* | *Epilan* | P | giant grouper |
| *Gobionotothen gibberifrons* | *Gobgib* | C | humped rockcod |
| *Gymnodraco acuticeps* | *Gymacu* | C | ploughfish |
| *Harpagifer antarcticus* | *Harant* | C | Antarctic spiny plunderfish |
| *Notothenia angustata* | *Notang* | C | Maori chief |
| *Notothenia coriiceps* | *Notcor* | C | black rockcod |
| *Nototheniops nudifrons* | *Notnud* | C | yellowfin notie |
| *Pagetopsis macropterus* | *Pagmac* | C | NA |
| *Pseudochaenichthys georgianus* | *Psegeo* | C | South Georgia icefish |
| *Psilodraco breviceps* | *Psibre* | C | NA |
| *Trematomus bernacchii* | *Treber* | C | emerald rockcod |
| *Trematomus borchgrevinki* | *Trebor* | C | bald notothen |
| *Trematomus loennbergii* | *Treloe* | C | Scaly rockcod |
| *Trematomus newnesi* | *Trenew* | C | Dusky rockcod |

[2] Groups: C = cryonotothenioid ice fish; N = non-ice fish notothenioid; P = perciform outgroup; T = teleost outgroup
[3] Common names from FishBase (https://www.fishbase.se/home.php)

### Table S3. Opsin sequence identifiers include in Figures 1 and S1-S5

| Species | Rh1[4] | Rh2 | SWS1 | SWS2 | LWS | References |
|---|---|---|---|---|---|---|
| *Anaoce* | XP_031700525 | XP_031708683<br>XP_031698504 | XP_031696717 | XP_031728565<br>XP_031728434 | XP_033482814 | |
| *Chaace* | HQ170035<br>HQ170036 | | | | | |
| *Chawil* | HQ170037 | | | | | |
| *Chaeso* | HQ170040 | | | | | |
| *Chagun* | HQ170041 | | BAJ15894* | | BAL15673** | *Miyazaki et al 2011;<br>**Miyazaki and Iwami, 2012 |
| *Chidew* | HQ170045 | | | | | |
| *Chiham* | HQ170047 | | | | | |
| *Chimye* | HQ170048<br>(HQ170049) | | | | | |
| *Chiras* | HQ170050<br>(HQ170051) | | | | | |
| *Cotgob* | XP_029291931* | XP_029286851*<br>XP_029286846* | XP_029282297* | XP_029288360*<br>XP_029288359* | XP_029287453* | *Bista et al 2020 |
| *Danrer* | BAC21668**<br>ADP06864*** | BAC24129*<br>BAC24130*<br>BAC24131*<br>BAC24132* | BAC24134* | BAC24133* | BAC24127*<br>BAC24128* | *Chinen et al 2002;<br>**Hamaoka et al 2002;<br>***Morrow et al. 2017 |
| *Dismaw* | DQ498794*** | KAF3856251*<br>AAV53958** | AAX99140** | | | *Jae Lee et al 2021<br>**Pointer et al. 2005<br>***Sanchez et al. 2007 |
| *Dollon* | HQ170034 | | | | | |
| *Elemac* | AY141303*** | evm.model.<br>scaffold423.2* | evm.model.<br>scaffold79.26*<br>(partial) | AJD38779**<br>AJD38778** | evm.model.<br>scaffold383.27*<br>(pseudogene?) | *Chen et al 2019<br>**Cortesi et al<br>***Sanchez et al. 2007 |
| *Epilan* | XP_033484724* | XP_033483237*<br>XP_033483238*<br>XP_033483239* | XP_033470999* | XP_033482114*<br>XP_033482113*<br>XP_033483640* | XP_033482814* | *Wang et al 2019 |
| *Gobgib* | | | | | BAL15677* | *Miyazaki and Iwami, 2012 |
| *Gymacu* | XP_034066827* | XP_034075023*<br>XP_034075029*<br>AAV53961** | XP_034055527*<br>AAX99145** | XP_034068225* | XP_034068146* | *NCBI Genome Annotation Release 100<br>**Pointer et al. 2005 |
| *Harant* | HQ170033 | | | | | |
| *Notang* | DQ498787** | AAV53960* | AAX99143* | | | *Pointer et al. 2005<br>**Sanchez et al. 2007 |
| *Notcor* | XP_010779059* | XP_010780753* | | XP_010769741*<br>(pseudogene?) | XP_010769740* | *Lin et al, 2017 and Shin et al, 2014 |
| *Notnud* | HM166262 | | | | BAL15676*<br>BAL15680* | *Miyazaki and Iwami, 2012 |
| *Pagmac* | EU637990*<br>(HQ170061) | | AAX99144** | | | *Li et al, 2009<br>**Pointer et al. 2005 |
| *Psegeo* | XP_033942519* | XP_033938873*<br>XP_033938753* | XP_033935414* | XP_033939966* | XP_033938770* | *NCBI Genome Annotation Release 100 |
| *Psibre* | AMW90782<br>(KU647484) | | | | | |
| *Treber* | XP_033982463* | XP_033994711*<br>XP_033994756* | XP_033973103* | XP_033977579*<br>XP_033977564* | XP_033977647* | *NCBI Genome Annotation Release 100 |
| *Trebor* | HM166264 | AAV53959* | AAX99142* | | | *Pointer et al. 2005 |
| *Tremloe* | HM166275<br>DQ498802** | | AAX99141* | AAV53962* | | *Pointer et al. 2005<br>**Sanchez et al. 2007 |
| *Trenew* | HM166276 | | | | BAL15675* | *Miyazaki and Iwami, 2012 |

[4] Sequences that encode identical proteins are listed in parentheses and excluded from analyses.

**Table S4. Changes in major tuning sites and their predicted $\lambda_{max}$[5]**

| | Rh1[6] | | | Rh2[7] | | SWS2[8] | SWS1[9] | | LWS[10] | |
|---|---|---|---|---|---|---|---|---|---|---|
| **Species** | **D83N ↓3 nm** | **E122Q ↓13 nm** | **A292S ↓2 nm** | **E122Q ↓13-17 nm** | **M207L ↓6-10 nm** | **W265Y ↓29 nm** | **S114A** | **S118A ↓5 nm** | **S164A ↓7 nm** | **S164P** |
| *Chaace* | 2/2 | 0/2 | 0/2 | 0/0 | 0/0 | 0/0 | 0/0 | 0/0 | 0/0 | 0/0 |
| *Chawil* | 1/1 | 0/1 | 0/1 | 0/0 | 0/0 | 0/0 | 0/0 | 0/0 | 0/0 | 0/0 |
| *Chaeso* | 0/1 | 0/1 | 0/1 | 0/0 | 0/0 | 0/0 | 0/0 | 0/0 | 0/0 | 0/0 |
| *Chagun* | 1/1 | 0/1 | 0/1 | 0/0 | 0/0 | 0/0 | 1/1 | 1/1 | 1/1 | 0/1 |
| *Chidew* | 1/1 | 0/1 | 1/1 | 0/0 | 0/0 | 0/0 | 0/0 | 0/0 | 0/0 | 0/0 |
| *Chiham* | 1/1 | 0/1 | 0/1 | 0/0 | 0/0 | 0/0 | 0/0 | 0/0 | 0/0 | 0/0 |
| *Chimye* | 1/1 | 0/1 | 0/1 | 0/0 | 0/0 | 0/0 | 0/0 | 0/0 | 0/0 | 0/0 |
| *Chiras* | 1/1 | 0/1 | 0/1 | 0/0 | 0/0 | 0/0 | 0/0 | 0/0 | 0/0 | 0/0 |
| *Dismaw* | 0/1 | 0/1 | 0/1 | 2/2 | 1/2 | 0/0 | 0/1 | 0/1 | 0/0 | 0/0 |
| *Dollon* | 0/1 | 0/1 | 1/1 | 0/0 | 0/0 | 0/0 | 0/0 | 0/0 | 0/0 | 0/0 |
| *Gobgib* | 0/0 | 0/0 | 0/0 | 0/0 | 0/0 | 0/0 | 0/0 | 0/0 | 1/1 | 0/1 |
| *Gymacu* | 1/1 | 0/1 | 1/1 | 3/3 | 3/3 | 0/1 | 2/2 | 0/2 | 1/1 | 0/1 |
| *Harant* | 0/1 | 0/1 | 0/1 | 0/0 | 0/0 | 0/0 | 0/0 | 0/0 | 0/0 | 0/0 |
| *Notang* | 0/1 | 0/1 | 0/1 | 0/1 | 0/1 | 0/0 | 1/1 | 1/1 | 0/0 | 0/0 |
| *Notcor* | 0/1 | 1/1 | 0/1 | 1/1 | 0/1 | 0/1 | 0/0 | 0/0 | 0/1 | 0/1 |
| *Notnud* | 0/1 | 1/1 | 0/1 | 0/0 | 0/0 | 0/0 | 0/0 | 0/0 | 2/2 | 0/2 |
| *Pagmac* | 1/1 | 0/1 | 0/1 | 0/0 | 0/0 | 0/0 | 0/1 | 0/1 | 0/0 | 0/0 |
| *Psegeo* | 1/1 | 0/1 | 0/1 | 2/2 | 2/2 | 1/1 | 0/1 | 0/1 | 1/1 | 0/1 |
| *Psibre* | 1/1 | 0/1 | 0/1 | 0/0 | 0/0 | 0/0 | 0/0 | 0/0 | 0/0 | 0/0 |
| *Treber* | 0/1 | 0/1 | 0/1 | 2/2 | 2/2 | 0/0 | 0/1 | 0/1 | 0/1 | 1/1 |
| *Trebor* | 0/1 | 0/1 | 0/1 | 1/1 | 1/1 | 1/2 | 0/1 | 0/1 | 0/0 | 0/0 |
| *Treloe* | 0/2 | 0/2 | 021 | 0/0 | 0/0 | 0/1 | 1/1 | 0/1 | 0/0 | 0/0 |
| *Trenew* | 0/1 | 0/1 | 0/1 | 0/0 | 0/0 | 0/0 | 0/0 | 0/0 | 1/1 | 0/1 |

5 Numbers indicate the number of sequences with indicated tuning site change / total number of sequences identified. Blue shading indicates predicted decreases in $\lambda_{max}$ and orange shading indicates an unknown impact on $\lambda_{max}$. Gray shading indicates that no sequences were identified.

6 Predicted shifts in $\lambda_{max}$ of Rh1 from Yokoyama et al (2008a). Note that the combination of D83N and A292S can decrease $\lambda_{max}$ by 14-17 nm. Changes in other major tuning site F261 (Lin et al 2017) were not observed.

7 Predicted shifts in $\lambda_{max}$ of Rh2 from Yokoyama et al (1999), Takenaka and Yokoyama (2007) and Yokoyama and Jia (2020). Changes in other major tuning sites S97 and A292 (Lin et al 2017) were not observed.

8 Predicted shifts in $\lambda_{max}$ of SWS2 from Yokoyama et al (2007). Changes in other major tuning sites A94, S97, and T118, (Lin et al 2017) were not observed.

9 Shi and Yokoyama (2003) demonstrated the reverse mutation A118S increases $\lambda_{max}$ of SWS1 by 5 nm, thus we predict a comparable decrease for S118A. Changes in other major tuning sites T52, F86, and Q93 (Lin et al 2017) were not observed.

10 Predicted shift for LWS S164A from Yokoyama et al (2008b). Changes in other major tuning sites H181, Y261, T269, and A292 (Lin et al 2017) were not observed.

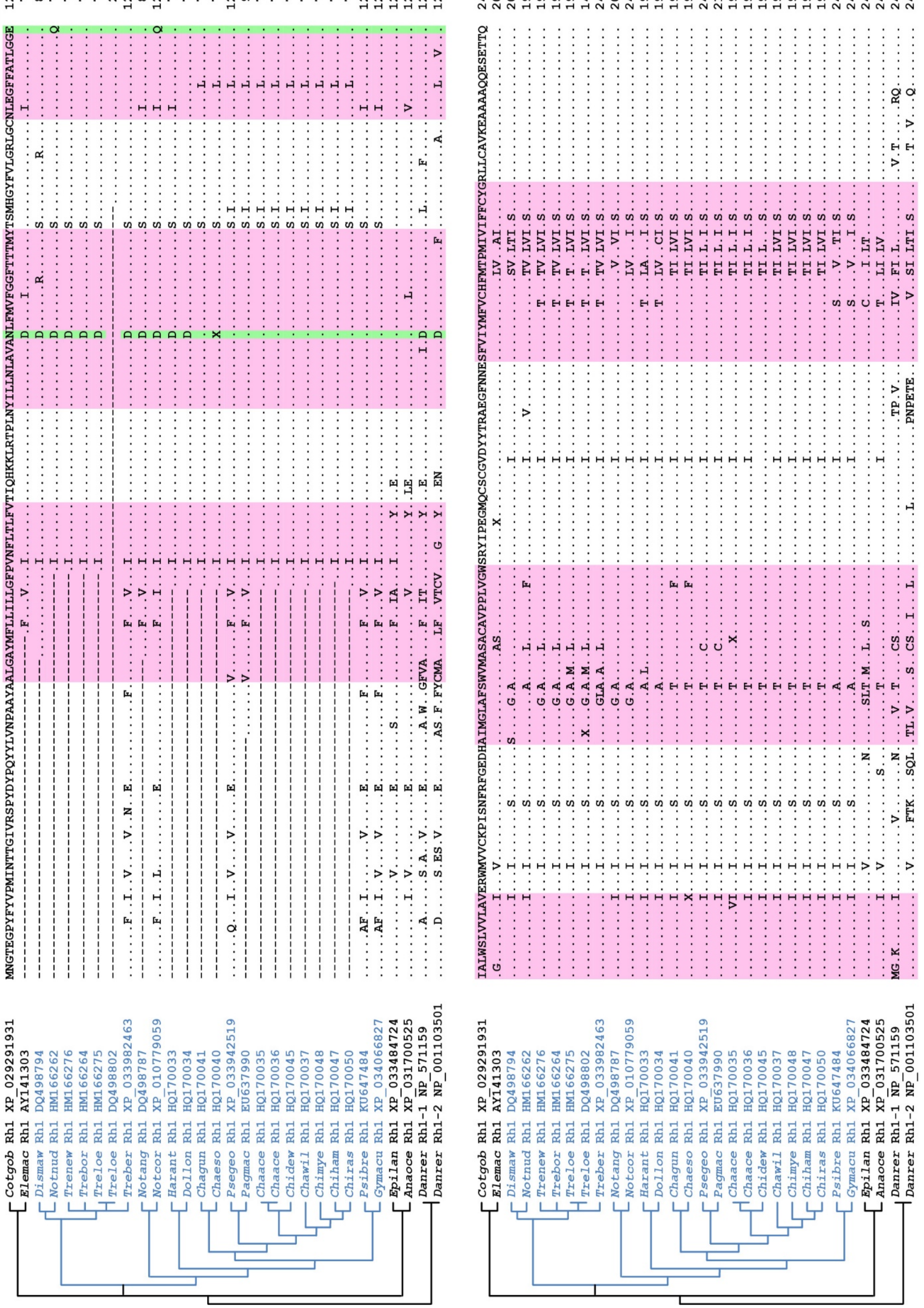

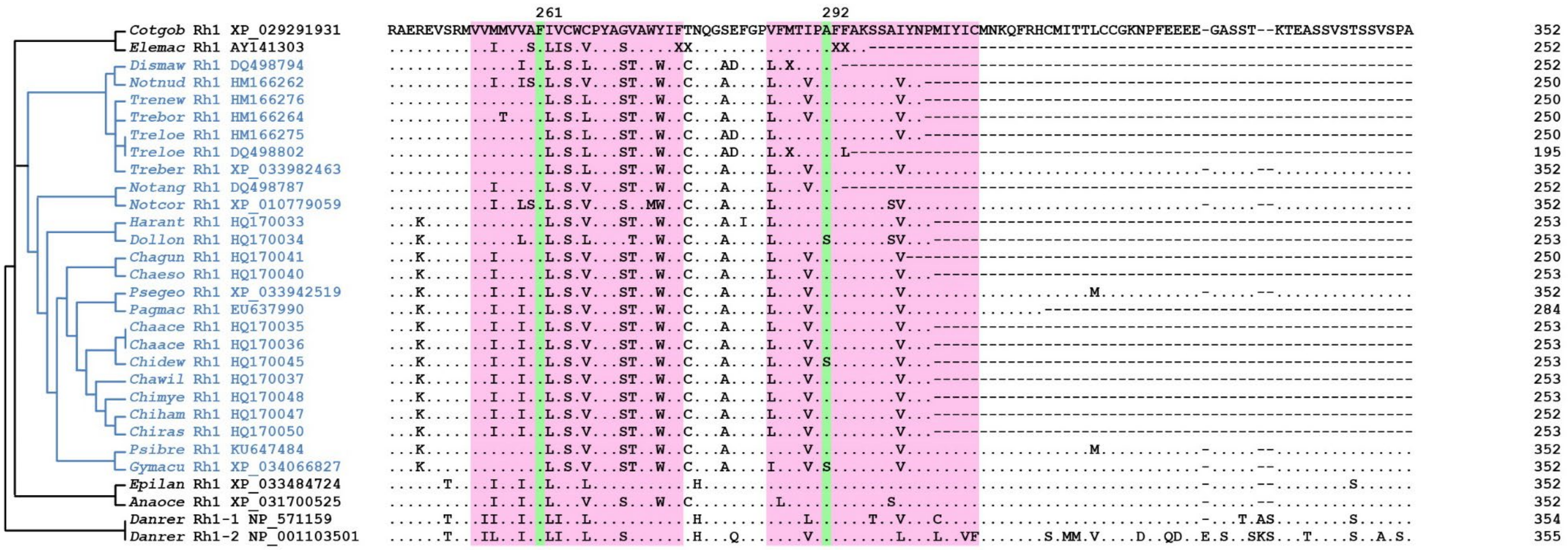

**Figure S1. Notothenioid Rh1 sequences and tuning sites**.

Alignment of representative cryonotothenioid Rh1 protein sequences compared with representative non-icefish notothenioid (*Cottoperca gobio, Cotgob and Eleginops maclovinus, Elemac*), perciforme (*Anarrhichthys ocellatus*, *Anaoce* and *Epinephelus lanceolatus, Epilan*) and zebrafish (*Danio rerio, Danrer*) Rh1 sequences. The phylogenetic trees (left) show the relationship between icefish (blue text), other perciformes, and zebrafish. The *Cotgob* sequence was used as a reference (top row) and dots in the alignment indicate identity. Differences from *Cotgob* are indicated by the appropriate amino acid symbol. Amino acids in transmembrane domains are highlighted pink. Major tuning site amino acids (Lin et al., 2017) are highlighted green with the position number of the amino acid at the top of each column. Position numbers are based on the bovine rhodopsin (Palczewski et al, 2000). Sequence identifiers are included in the alignment and listed in Supplemental Table S2. Genus and species are included in Supplemental Table S1.

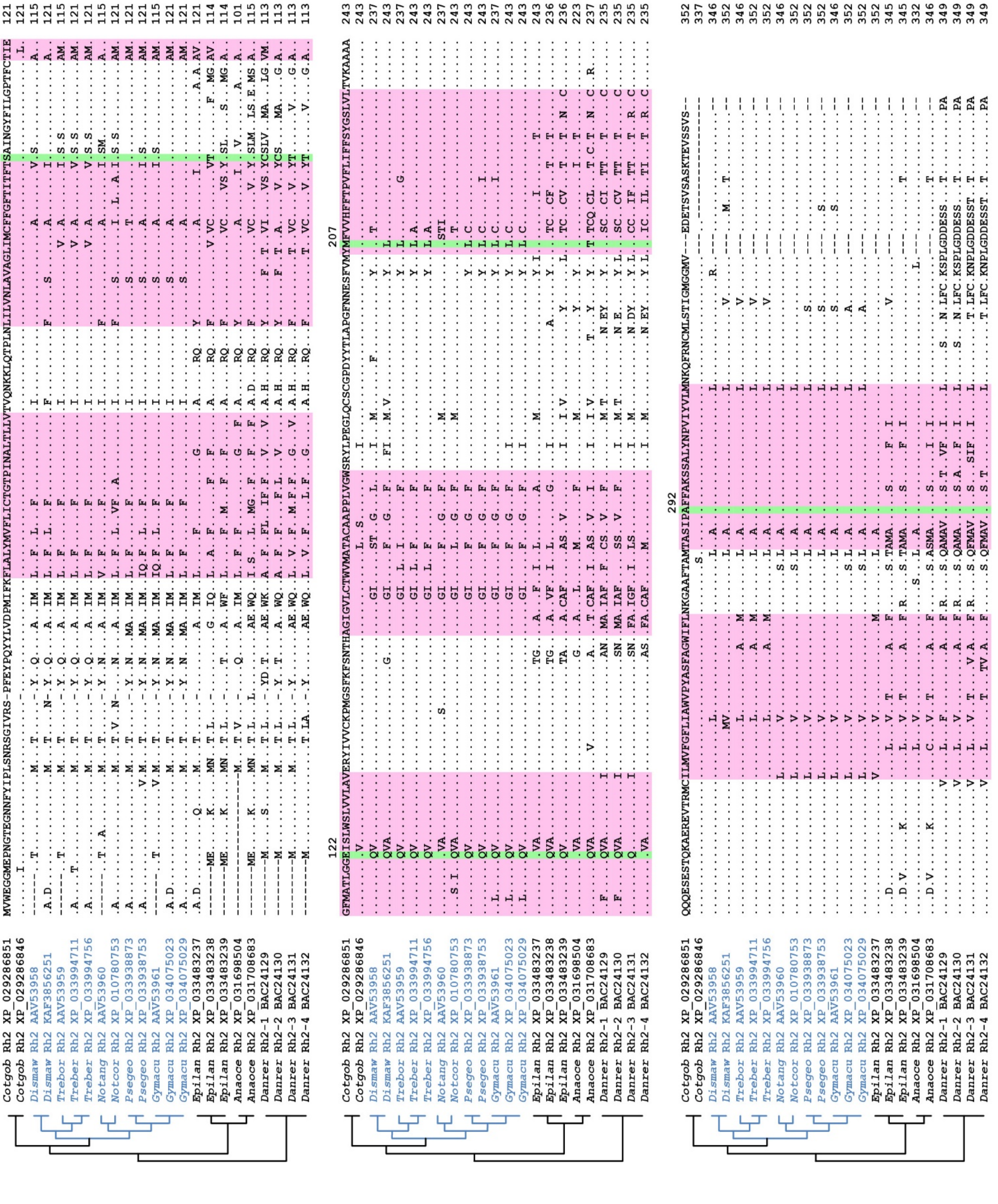

**Figure S2. Notothenioid Rh2 sequences and tuning sites.**
Alignment of representative cryonotothenioid Rh2 protein sequences compared with representative non-icefish notothenioid (*Cottoperca gobio, Cotgob*), perciforme (*Anarrhichthys ocellatus, Anaoce* and *Epinephelus lanceolatus, Epilan*) and zebrafish (*Danio rerio, Danrer*) Rh2 sequences. The phylogenetic trees (left) show the relationship between icefish (blue text), other perciformes, and zebrafish. A *Cotgob* sequence was used as a reference (top row) and dots in the alignment indicate identity. Differences from *Cotgob* are indicated by the appropriate amino acid symbol. Amino acids in transmembrane domains are highlighted pink. Major tuning site amino acids (Lin et al., 2017) are highlighted green with the position number of the amino acid at the top of each column. Position numbers are based on bovine rhodopsin (Palczewski et al, 2000). Sequence identifiers are included in the alignment and listed in Supplemental Table S2. Genus and species are included in Supplemental Table S1.

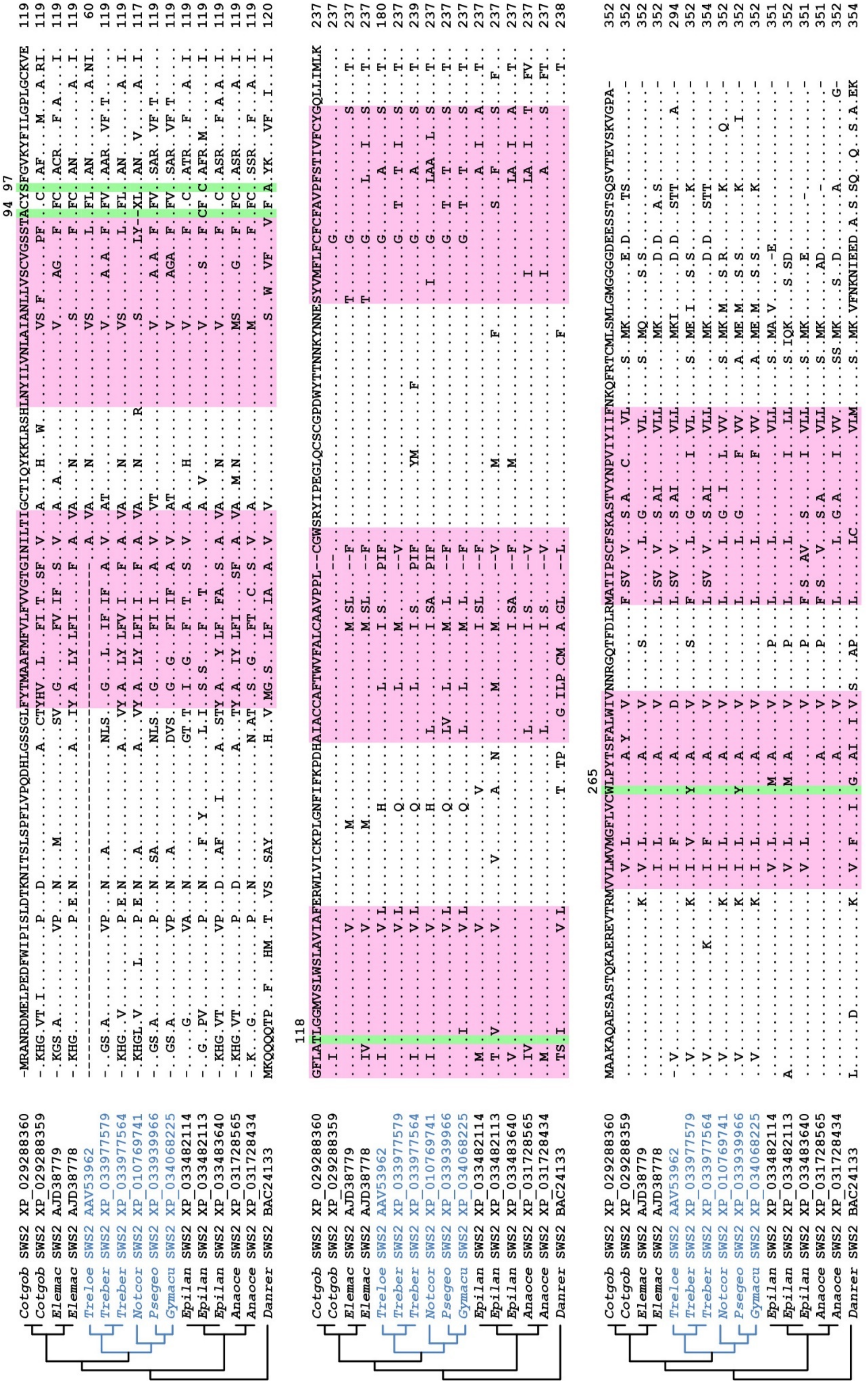
Cotgob SWS2 XP_029288360
Cotgob SWS2 XP_029288359
Elemac SWS2 AJD38779
Elemac SWS2 AJD38778
Treloe SWS2 AAV53962
Treber SWS2 XP_033977579
Treber SWS2 XP_033977564
Notcor SWS2 XP_010769741
Psegeo SWS2 XP_033939966
Gymacu SWS2 XP_034068225
Epilan SWS2 XP_033482114
Epilan SWS2 XP_033482113
Epilan SWS2 XP_033483640
Anaoce SWS2 XP_031728565
Anaoce SWS2 XP_031728434
Danrer SWS2 BAC24133
94 97
118
265

**Figure S3. Notothenioid SWS2 sequences and tuning sites.**
Alignment of representative cryonotothenioid SWS2 protein sequences compared with representative non-icefish notothenioid (*Cottoperca gobio, Cotgob* and *Eleginops maclovinus, Elemac*), perciforme (*Anarrhichthys ocellatus, Anaoce* and *Epinephelus lanceolatus, Epilan*) and zebrafish (*Danio rerio, Danrer*) SWS2 sequences. The phylogenetic trees (left) show the relationship between icefish (blue text), other perciformes, and zebrafish. A *Cotgob* sequence was used as a reference (top row) and dots in the alignment indicate identity. Differences from *Cotgob* are indicated by the appropriate amino acid symbol. Amino acids in transmembrane domains are highlighted pink. Major tuning site amino acids (Lin et al., 2017) are highlighted green with the position number of the amino acid at the top of each column. Position numbers are based on bovine rhodopsin (Palczewski et al, 2000). Sequence identifiers are included in the alignment and listed in Supplemental Table S2. Genus and species are included in Supplemental Table S1.

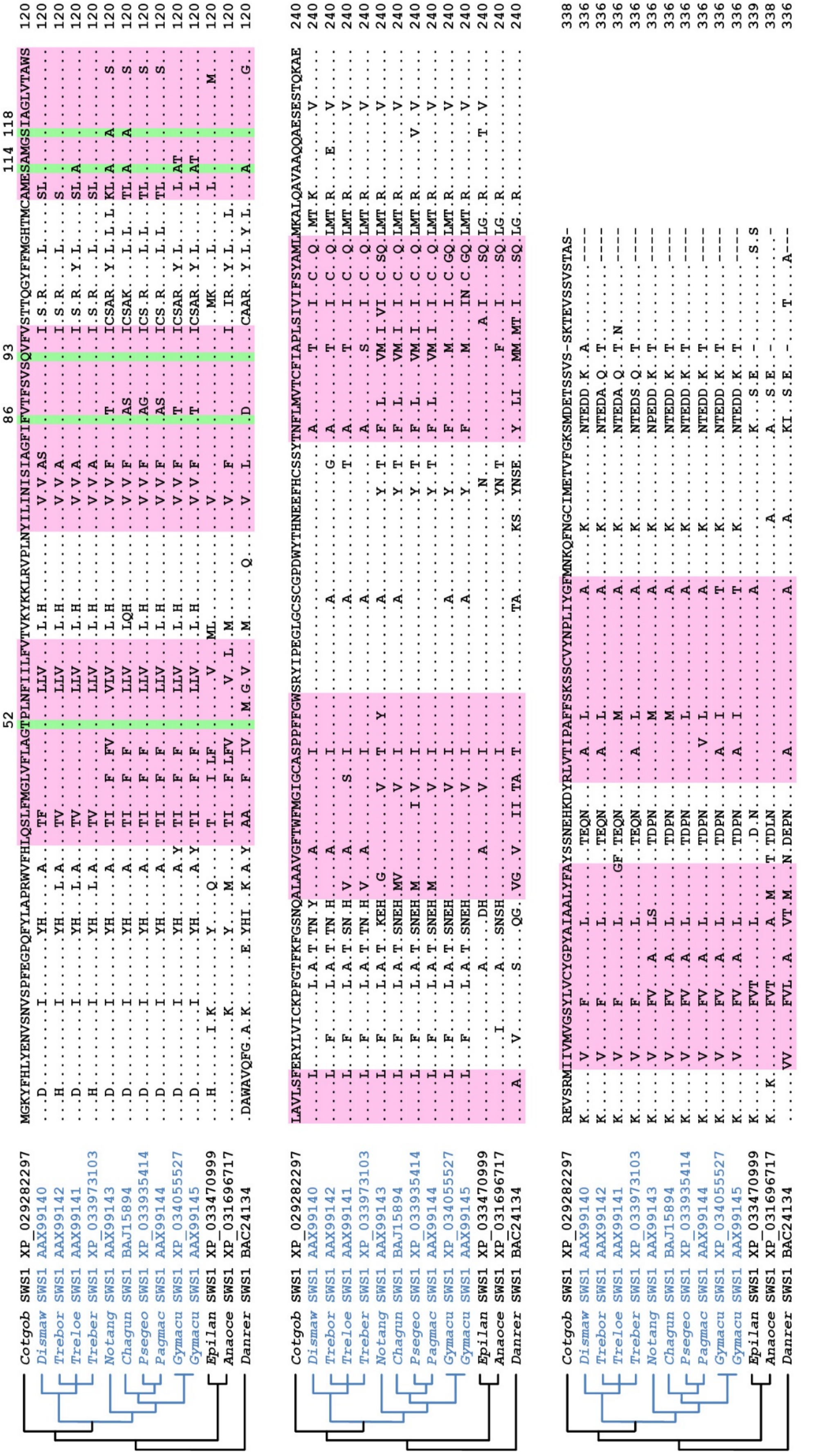

**Figure S4. Notothenioid SWS1 sequences and tuning sites.**
Alignment of representative cryonotothenioid SWS1 protein sequences compared with representative non-icefish notothenioid (*Cottoperca gobio, Cotgob*), perciforme (*Anarrhichthys ocellatus*, *Anaoce* and *Epinephelus lanceolatus, Epilan*) and zebrafish (*Danio rerio, Danrer*) SWS1 sequences. The phylogenetic trees (left) show the relationship between icefish (blue text), other perciformes, and zebrafish. The *Cotgob* sequence was used as a reference (top row) and dots in the alignment indicate identity. Differences from *Cotgob* are indicated by the appropriate amino acid symbol. Amino acids in transmembrane domains are highlighted pink. Major tuning site amino acids (Lin et al., 2017) are highlighted green with the position number of the amino acid at the top of each column. Position numbers are based on bovine rhodopsin (Palczewski et al, 2000). Sequence identifiers are included in the alignment and listed in Supplemental Table S2. Genus and species are included in Supplemental Table S1.

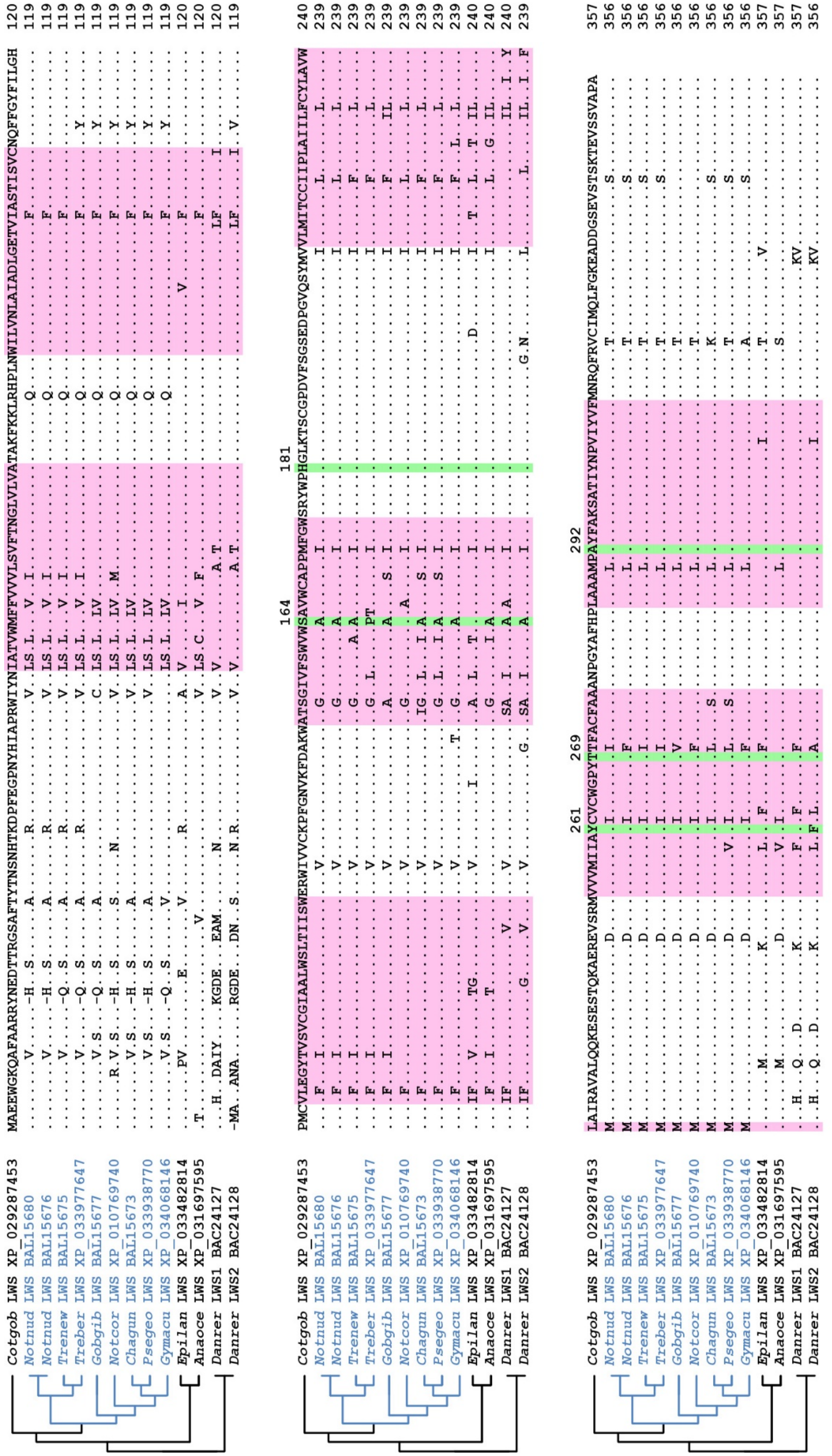

**Figure S5. Notothenioid LWS sequences and tuning sites.**
Alignment of representative cryonotothenioid LWS protein sequences compared with representative non-icefish notothenioid (*Cottoperca gobio, Cotgob*), perciforme (*Anarrhichthys ocellatus*, *Anaoce* and *Epinephelus lanceolatus, Epilan*) and zebrafish (*Danio rerio, Danrer*) LWS sequences. The phylogenetic trees (left) show the relationship between icefish (blue text), other perciformes, and zebrafish. The *Cotgob* sequence was used as a reference (top row) and dots in the alignment indicate identity. Differences from *Cotgob* are indicated by the appropriate amino acid symbol. Amino acids in transmembrane domains are highlighted pink. Major tuning site amino acids (Lin et al., 2017) are highlighted green with the position number of the amino acid at the top of each column. Position numbers are based on bovine rhodopsin (Palczewski et al, 2000). Sequence identifiers are included in the alignment and listed in Supplemental Table S2. Genus and species are included in Supplemental Table S1.

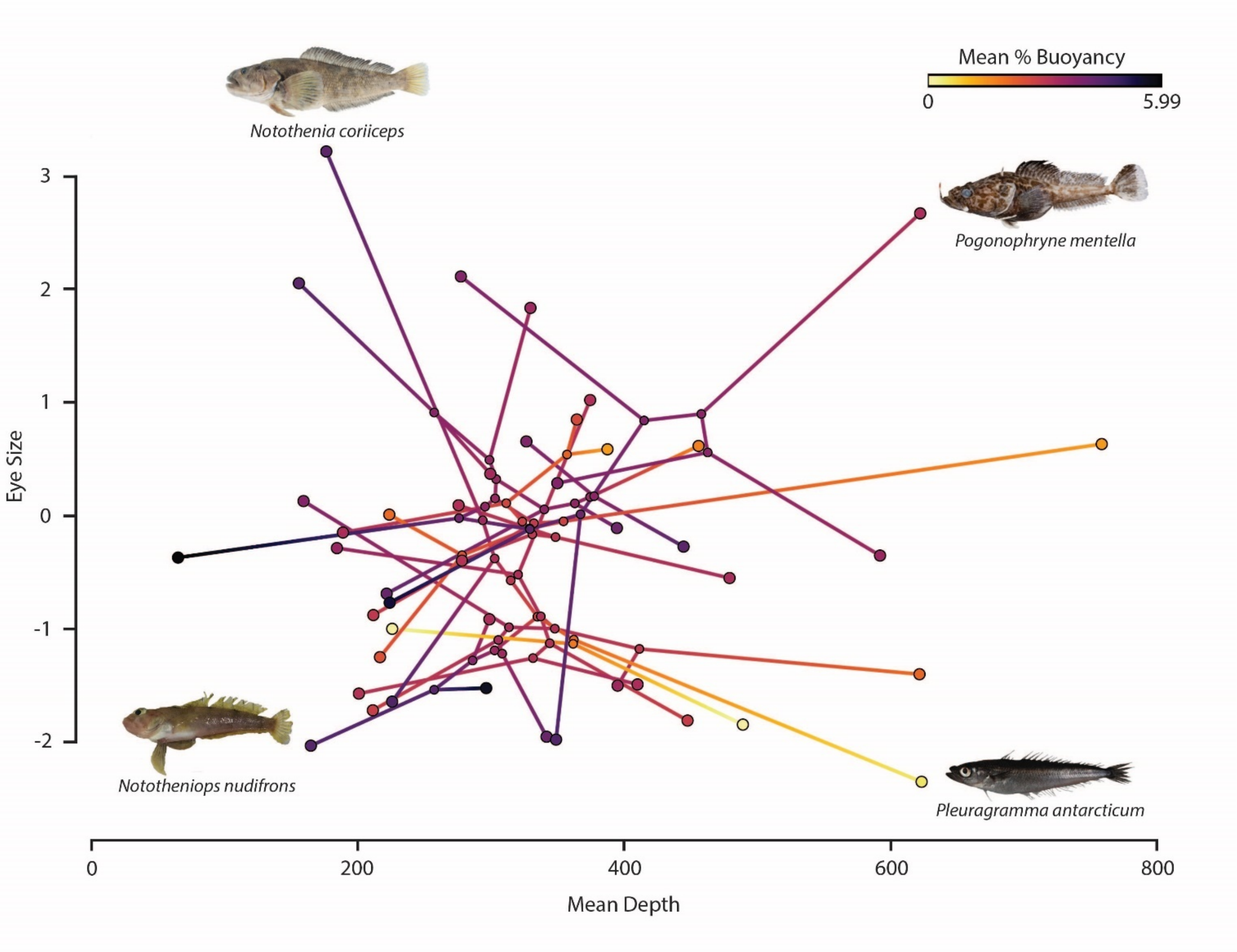

**Figure S6. Phylomorphospace plot of depth versus eye size.**
Phylomorphospace depicting projection of notothenioid phylogeny into trait space defined by mean depth of occurrence on the X axis and relative eye size on the Y axis. Larger shaded circles correspond to each of the notothenioid species sampled in our phylogeny, while smaller shaded circles correspond to nodes in the phylogeny. The ancestral state reconstruction of mean percentage buoyancy (% B) has been mapped onto the projected phylogeny to facilitate simultaneous visualization of evolutionary changes in mean depth of occurrence, eye size, and buoyancy.

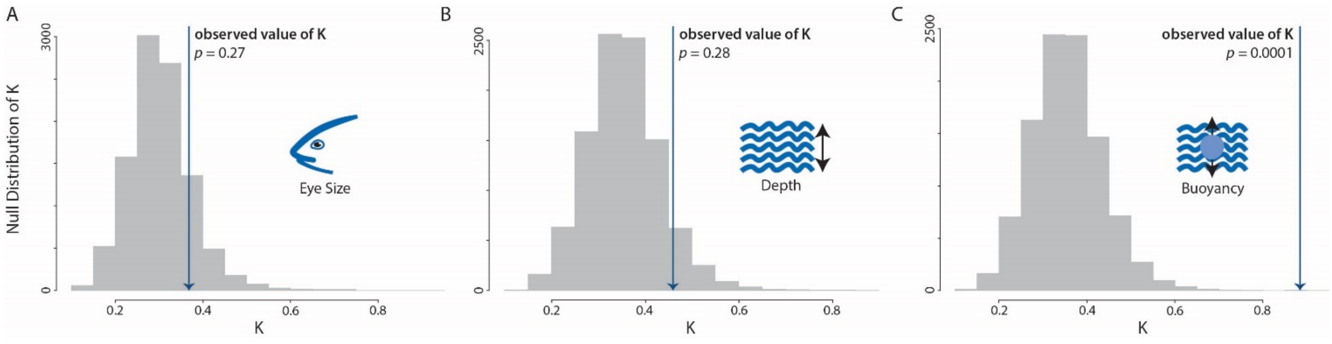

**Figure S7. Phylogenetic signal in ecomorphological traits.**
Comparisons of empirical values of Blomberg et al.'s K (2003) calculated for relative eye size (A), mean depth of occurrence (B), and mean percentage buoyancy (C) to distribution of K values calculated from 10,000 permutations of trait evolution on the notothenioid phylogeny.

**References**.

Bista I, McCarthy SA, Wood J, Ning Z, Detrich Iii HW, Desvignes T, Postlethwait J, Chow W, Howe K, Torrance J, Smith M, Oliver K; Vertebrate Genomes Project Consortium, Miska EA, Durbin R. 2020. The genome sequence of the channel bull blenny, Cottoperca gobio (Günther, 1861). *Wellcome Open Res*. 24; 5:148.

Blomberg SP, Garland T, Ives AR. 2003. Testing for phylogenetic signal in comparative data: behavioral traits are more labile. Evolution. 57(4): 717-45.

Chen L, Lu Y, Li W, Ren Y, Yu M, Jiang S, Fu Y, Wang J, Peng S, Bilyk KT, Murphy KR, Zhuang X, Hune M, Zhai W, Wang W, Xu Q, Cheng CC. 2019. The genomic basis for colonizing the freezing Southern Ocean revealed by Antarctic toothfish and Patagonian robalo genomes. *Gigascience*. 8(4): giz016.

Chinen A, Hamaoka T, Yamada Y, Kawamura S. 2003. Gene duplication and spectral diversification of cone visual pigments of zebrafish. *Genetics*. 163(2): 663-75.

Cortesi F, Musilová Z, Stieb SM, Hart NS, Siebeck UE, Malmstrøm M, Tørresen OK, Jentoft S, Cheney KL, Marshall NJ, Carleton KL, Salzburger W. 2015. Ancestral duplications and highly dynamic opsin gene evolution in percomorph fishes. *Proc Natl Acad Sci U S A*.112(5):1493-8.

Fujiyabu C, Sato K, Utari NML, Ohuchi H, Shichida Y, Yamashita T. 2019. Evolutionary history of teleost intron-containing and intron-less rhodopsin genes. *Sci Rep*. 9(1): 10653.

Hamaoka T, Takechi M, Chinen A, Nishiwaki Y, Kawamura S. 2002. Visualization of rod photoreceptor development using GFP-transgenic zebrafish. *Genesis*. 34(3): 215-20.

Jae Lee S, Kim JH, Jo E, Choi E, Kim J, Choi SG, Chung S, Kim HW, Park H. 2021. Chromosomal assembly of the Antarctic toothfish (Dissostichus mawsoni) genome using third-generation DNA sequencing and Hi-C technology. *Zool Res*. 42(1): 124-129.

Li B, Dettaï A, Cruaud C, Couloux A, Desoutter-Meniger M, Lecointre G. 2009. RNF213, a new nuclear marker for acanthomorph phylogeny. *Mol Phylogenet Evol*. 50(2): 345-63.

Lin JJ, Wang FY, Li WH, Wang TY. 2017. The rises and falls of opsin genes in 59 ray-finned fish genomes and their implications for environmental adaptation. *Sci Rep*. 7(1): 15568.

Miyazaki T, Iwami T. 2012. Molecular cloning of cDNA encoding red opsin gene in the retinas of five Antarctic notothenioid fishes. *Polar Biol*. 35: 775-783.

Miyazaki T, Nakata M, Kasagi S, Iwami T, Yamauchi M, Kawamura S. 2011. Molecular cloning of ultraviolet-sensitive visual pigment in juvenile Champsocephalus gunnari (Channichthyidae). *Polar Biol*. 34: 235-242.

Morrow JM, Lazic S, Dixon Fox M, Kuo C, Schott RK, de A Gutierrez E, Santini F, Tropepe V, Chang BS. 2017. A second visual rhodopsin gene, rh1-2, is expressed in zebrafish photoreceptors and found in other ray-finned fishes. *J Exp Biol*. 220(Pt 2): 294-303.

Palczewski K, Kumasaka T, Hori T, Behnke CA, Motoshima H, Fox BA, Le Trong I, Teller DC, Okada T, Stenkamp RE, Yamamoto M, Miyano M. 2000. Crystal structure of rhodopsin: A G protein-coupled receptor. *Science*. 289(5480): 739-45.

Pointer MA, Cheng CH, Bowmaker JK, Parry JW, Soto N, Jeffery G, Cowing JA, Hunt DM. 2005. Adaptations to an extreme environment: retinal organisation and spectral properties of photoreceptors in Antarctic notothenioid fish. *J Exp Biol*. 208(Pt 12): 2363-76.

Sabaj MH. 2016. Standard symbolic codes for institutional resource collections in herpetology and ichthyology: an Online Reference. Version 6.5 (16 August 2016). Electronically accessible at http://www.asih.org/, American Society of Ichthyologists and Herpetologists, Washington, DC.

Sanchez S, Dettai A, Bonillo C, Ozouf-Costaz C, Detrich III W, Lecointre G. 2007. Molecular and morphological phylogenies of the Antarctic teleostean family Nototheniidae, with emphasis on the Trematominae. *Polar Biol*. 30: 155-166.

Shin SC, Ahn DH, Kim SJ, Pyo CW, Lee H, Kim MK, Lee J, Lee JE, Detrich HW, Postlethwait JH, Edwards D, Lee SG, Lee JH, Park H. 2014. The genome sequence of the Antarctic bullhead notothen reveals evolutionary adaptations to a cold environment. *Genome Biol*. 15(9): 468.

Takenaka N, Yokoyama S. 2007. Mechanisms of spectral tuning in the RH2 pigments of Tokay gecko and American chameleon. *Gene*. 399(1): 26-32.

Wang D, Chen X, Zhang X, Li J, Yi Y, Bian C, Shi Q, Lin H, Li S, Zhang Y, You X. 2019. Whole Genome Sequencing of the Giant Grouper (Epinephelus lanceolatus) and High-Throughput Screening of Putative Antimicrobial Peptide Genes. *Mar Drugs*. 17(9): 503.

Yokoyama S, Jia H. 2020. Origin and adaptation of green-sensitive (RH2) pigments in vertebrates. FEBS Open Bio. 10(5): 873-882.

Yokoyama S, Tada T, Zhang H, Britt L. 2008a. Elucidation of phenotypic adaptations: Molecular analyses of dim-light vision proteins in vertebrates. *Proc Natl Acad Sci U S A*. 105(36): 13480-5.

Yokoyama S, Takenaka N, Blow N. 2007. A novel spectral tuning in the short wavelength-sensitive (SWS1 and SWS2) pigments of bluefin killifish (*Lucania goodei*). *Gene*. 396(1): 196-202.

Yokoyama S, Yang H, Starmer WT. 2008b. Molecular basis of spectral tuning in the red- and green-sensitive (M/LWS) pigments in vertebrates. *Genetics*. 179(4): 2037-43.

Yokoyama S, Zhang H, Radlwimmer FB, Blow NS. 1999. Adaptive evolution of color vision of the Comoran coelacanth (*Latimeria chalumnae*). *Proc Natl Acad Sci U S A*. 96(11): 6279-84.
